# A probabilistic model to identify the core microbial community

**DOI:** 10.1101/491183

**Authors:** Thiago Gumiere, Kyle M. Meyer, Adam R. Burns, Silvio J. Gumiere, Brendan J. M. Bohannan, Fernando D. Andreote

## Abstract

The core microbial community has been hypothesized to have essential functions ranging from maintaining health in animals to protection against plant disease. However, the identification of the core microbial community is frequently based on arbitrary thresholds, selecting only the most abundant microorganisms. Here, we developed and tested an approach to identify the core community based on a probabilistic model. The Poisson distribution was used to identify OTUs with a probable occurrence in every sample of a given dataset. We identified the core communities of four extensive microbial datasets, and compared the results with conventional, but arbitrary, methods. The datasets were composed of the microbiomes of humans (tongue, gut, and skin), mice (gut), plant (grapevine) tissue, and the maize rhizosphere. Our proposed method revealed core microbial communities with higher richness and diversity than those previously described. This method also includes a greater number of rare taxa in the core, which are often neglected by arbitrary threshold methods. We demonstrated that our proposed method revels a probable core microbial community for each different habitat, which extend our knowledge about shared microbial communities. Our proposed method may help the next steps proving the essential functions of core microbial communities.

**Originality-Signifìcance Statement:** More rigorous and less arbitrary statistical methods could increase knowledge regarding the role of microorganisms and their interactions. Here, we suggest a probabilistic method to identify the microbial core community across systems. Our method identifies a large proportion of the rare community that likely belongs to the microbial core community, which was not identified by conventional methods. Our probabilistic model is a non-arbitrary approach to defining the microbial core community, which may help in the next step of the microbial core community studies.

## INTRODUCTION

The composition of microbial communities can vary greatly even over fine spatial and temporal scales, making it difficult to identify the drivers of community dynamics and the link between composition and function. To overcome the obfuscating effects of this variation, researchers often limit their focus to the ‘core’ community, which is defined as organisms that are ubiquitous in a given habitat, despite environmental fluctuation (Hamady and Knight, 2009). In microbial ecology, the core community refers to microbial taxa (Shade and Handelsman, 2012), or genes (Turnbaugh *et al*., 2007), shared across a set of samples in a given ecosystem.

There are considerable attempts to identify the core community across different hosts including corals (Ainsworth *et al*., 2015), zebrafish (Roeselers *et al*., 2011), mice (Pédron *et al*., 2012), ruminants (Henderson *et al*., 2015), *Arabidopsis thaliana* (Lundberg *et al*., 2012) and sugarcane plants (Yeoh *et al*., 2015). It has been suggested that the core microbial community could play essential roles in ecosystem functioning, and may also be useful as indicators of system perturbation (Shade and Handelsman, 2012; Saunders *et al*., 2015). For example, an abundant microbial core was identified across 210 human adult fecal samples, varying substantially in geographic origin, ethnic background and diet (Sekelja *et al*., 2011). The authors suggested that this core has an important role in gut homeostasis and health. Other studies have suggested roles for the core in plant growth promotion and the maintenance of plant health (Schlaeppi *et al*., 2014). However, few studies have been successful in directly linking the core microbial community to important community or ecosystem functions.

The lack of evidences for the importance of the core community may be due to how the core is identified. Since the core is defined to be ubiquitous in a habitat, it is assumed that the microbial taxa or genes belonging to the core should be found in every sample collected from a given habitat. The core microbial community is identified by identifying shared microorganisms or genes across a collection of samples *(discussed by* Shade and Handelsman, 2012). In this approach, the core is represented by taxa found in every sample analyzed (100% frequency across samples). However, to date no methodological approach has fully assessed the microbial diversity of any environmental sample (Kanagawa, 2003; Feinstein *et al*., 2009; Prosser, 2015). Current sequencing methods used to survey complex microbial communities tend to target the most abundant groups of microorganisms (Caporaso *et al*., 2011). Consequently, the rare component of the core microbial community is missed in these studies. The most commonly used approach to circumvent this problem is the definition of cutoffs for the frequency of microbes or genes to be classified as a member of the core microbial. For instance, researchers have used cutoff values ranging from 30% to 99% frequency across samples (Li *et al*., 2013; Ainsworth *et al*., 2015) to define the core community in environmental samples. However, these cutoffs still do not include rare taxa and also could result in false assignments to the core, thus influencing inferences about its function and composition.

Given the numerous difficulties associated with sampling and fully sequencing microbial communities, one solution to identify core community members is to use a probabilistic model to assign members of the microbial community to the core community. Here, we develop and test an approach to identifying the core community based on the Poisson distribution. Given the occurrence distribution of an event, *i.e*. a microorganism, in a group of samples, this model estimates the probability of this event in a group of samples (Rao and Rubin, 1964). Among discrete probability models, we selected the Poisson distribution because it is particularly suitable for large count datasets, *e.g*. a high number of events, and the occurrence of small or rare probabilities (Karlis, 2003), situations common when using microbial datasets to estimate a core community. Unlike other attempts to define the core community (e.g. Turnbaugh *et al*., 2009) there is no abundance threshold in our proposed method, which allows inclusion of rare taxa as possible members of the core microbial community.

We tested our proposed method using several previously published datasets, and compared our results to those obtained using conventional (i.e. arbitrary threshold) approaches. These datasets included human, mice, plant (grapevine tissue and maize rhizosphere), and soil data, and were obtained from the Earth Microbiome Project (EMP; http://www.earthmicrobiome.org). We hypothesized that our approach would lead to the identification of a probable core community that would be a higher proportion of the microbial community, and would also be composed of more microorganisms with low abundances (rare community members), than the core community identified using conventional approaches.

## RESULTS AND DISCUSSION

### Testing the distribution models and rarefaction effect

The first step was to select the most appropriate probabilistic method that fitted in OTU distributions. We tested 13 different models (described in Supplementary Material), and in Figure S1, we can observe the fourth best distribution models (Poisson, Chi-squared, Gamma and Beta) fitted on each dataset (Human, Grape, Maize and Mice). The Poisson distribution showed the higher and significant fit on OTU distribution, which is indicated by R^2^ and p-value < 0.05 in Table S1. We also observed that the Poisson distribution indicated lesser value of RMSE. Models based on ‘Poissonization’ arguments has also been indicated as good predictor of microbial unknown (Lladser *et al*., 2011).

The use of rarefaction, normalization method which equalizes the number of sequences (or reads) per sample, is discussed in the literature. According to McMurdie and Holmes (2014), the rarefaction increases the number of false positives species, and also with different abundance across sample classes. However, other simulation studies indicated that the rarefication is better than other normalization methods, clustering samples as biological origin (Weiss *et al*., 2017). As probability models requires the normalization, we evaluated the effect of the rarefaction on our proposed.

It can be observed in Figures S2, S3, S4 and S5 that the rarefaction method affects the line of Poisson distribution identification. We also observed that the values of R^2^ decreases with the increase of rarefication levels. However, the number of OTU’s identified as probable members of the core microbial community did not present a significant variation in general (Table S2). In grape dataset only the two highest rarefication levels, and in maize and human dataset only the lesser rarefication level showed a significant different number of core OTU’s identified. As indicated in Figures S6, S7, S8 and S9, the taxonomic composition at the phyla level was not significant affect by the most of rarefication levels. We verified the similarity of core community composition by different rarefication levels using NMDS analyses (Jaccard similarity). In Figure S10, we can observe that only the lowest level of rarefaction for the grape (Core_500), maize (Core_100), and human (core_100) datasets showed a significant difference from the other rarefaction levels. For the mice dataset, we observe the lower variation than the other datasets, but with the same pattern (lowest rarefaction level is not grouped). Considering this normalization effect, we decided to maintain the same method (rarefaction level) used by the authors of each published datasets for the next steps.

### A probabilistic method to identify the core microbial community

Using this probabilistic model, we identified core microbial communities for each dataset selected for analysis with R^2^ varying between 0.46 (mice) and 0.91 (grape), and with *p-values* as lower than 0.05. The obtained curves indicated the occurrence of OTUs with distinct values of frequency occurrence as components of the core microbial communities, which is not observed when other approaches are used (Figure 1 and Supplementary Figures S11, S12, and S13). As the results were based on a probabilistic method, we expected that our proposed method would identify a group closer to the real core community than the group identified by conventional methods.

**Figure 1.**
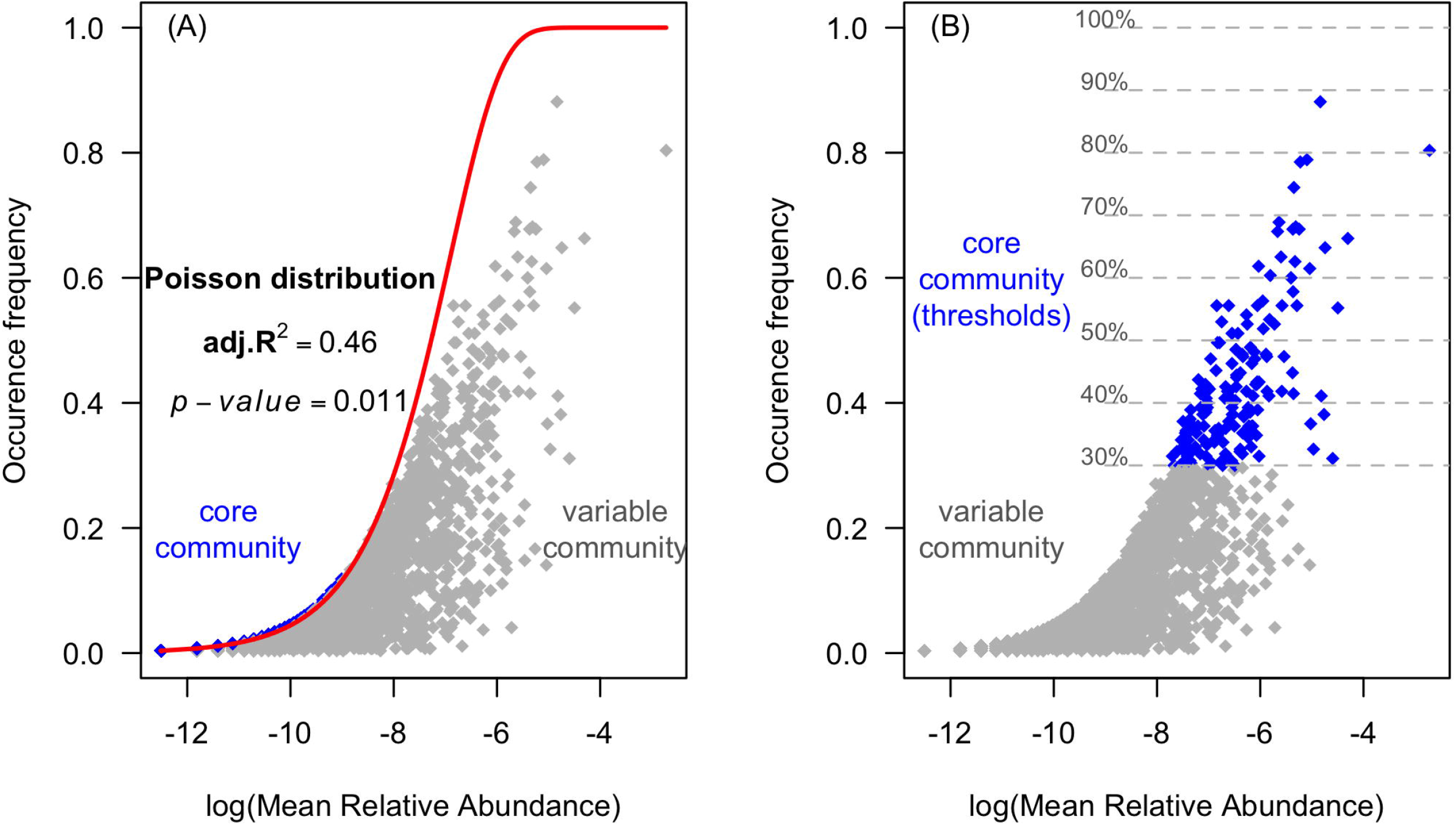
The core and variable communities of the mice microbiome determined by (A) our proposed method based on the Poisson distribution and (B) an arbitrary, threshold-based method.

We observed that our probabilistic method reveals a rich and diverse group of microorganism which has not been identified by conventional methods, but belong to the probable core microbial community. For example, the core microbial community identified in the mice database is composed of 170 OTUs using an arbitrary threshold of 30% detection frequency, and 1,717 OTUs using the method based on the Poisson distribution (Table 2). In particular, these differences were found for the occurrence of OTUs with low abundance, much more pronounced in the core community obtained by the method based on the Poisson distribution (e.g. Figure 1).

In the literature, the microorganisms with low abundance are frequently referred to as the “rare biosphere” (Sogin *et al*., 2006). The rare biosphere was first described as microorganisms with low growth rates, which could act as a “seed bank” of species or genes important in maintaining the functional redundancy of a system (Pedrós-Alió, 2006). These taxa could become dominant (in high abundance) under certain conditions (Shade *et al*., 2014). Following this view, members of the rare community can be classified as conditionally rare taxa (CRT), suggested to be ubiquitous in some systems (Shade and Gilbert, 2015). As members of a core microbial community, the CRT could be important to the stability and functional resilience of a system. Using our methodology, these groups could be properly classified within the core community, while the arbitrarily defined core rarely included these putative CRTs, likely due to their lower frequency (e.g. Figure 1B). The cut-offs for the core may fail to identify members of the core microbial community, *i.e*. this method may produce “false negatives”. By failing to include members in the core *(e.g*. low abundance taxa that are ubiquitous), researchers may be underestimating the contribution of the core to ecosystem function. Data from the mice dataset (Turnbaugh *et al*., (2009) did not identify a core microbial community across 100% of samples, or also using the PSM with abundance threshold. The probabilistic method identified the same three phyla as the arbitrary cutoff method *(Actinobacteria, Bacterioidetes*, and *Firmicutes)*, but also recovered an additional eight phyla *(Cyanobacteria, Fusobacteria, Lentisphaerae, Proteobacteria, Synergistetes, Tenericutes*, *TM7*, and *Verrucomicrobia*) as members of the core microbial community (Figure 2). The authors also indicated the distinct proportions of the *Bacteroidetes* and *Actinobacteria* phyla associated to obese and lean mice. Both phyla were also detected by our probabilistic method, with OTUs affiliated with these groups as components of the core microbial community.

**Figure 2.**
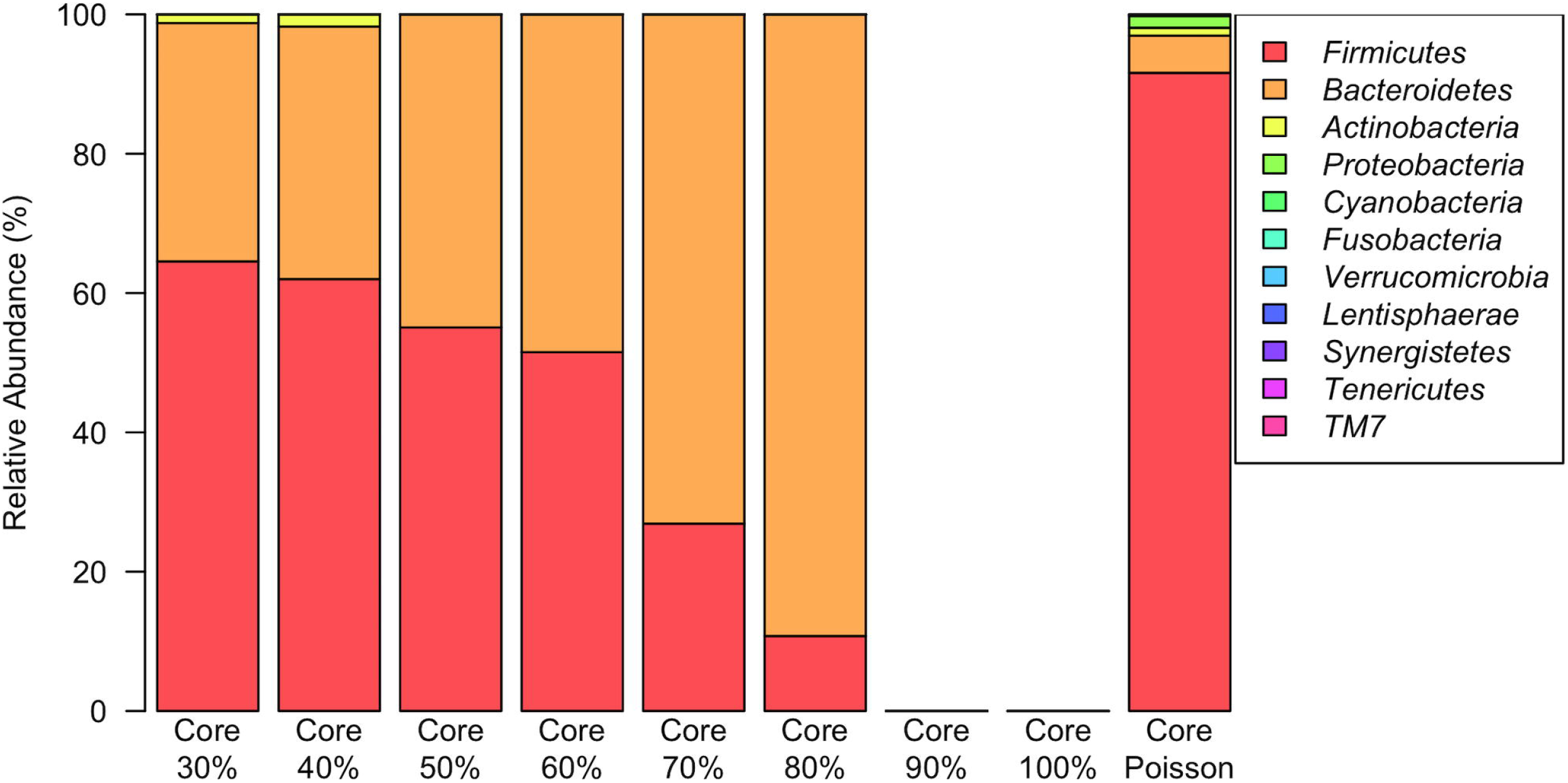
Percentage of the relative abundance of the core communities of the mice database determined by arbitrary methods (thresholds of 30,40,50,60,70,80,90 and 100%) and by our proposed method (Core Poisson).

Rather than defining a specific, core cutoffs, some researchers have used the term ‘persistent’ – referring to taxa with a high (but below 100%) occurrence frequency, or ‘transient’ referring to taxa with low occurrence frequency. For example, Caporaso *et al*., (2011) have identified a persistent and transient communities, which are classified as OTUs occurring in 60% or 20% of samples, respectively. Using this dataset (Caporaso *et al*. 2011), we identified a probable core community, also based on OTUs, across all of the human site samples made of 8,751 OTUs (Supplementary Figure S10). The authors identified classes belonging to the phyla *Firmicutes, Proteobacteria, Bacteroidetes*, and *Tenericutes* in the human gut. Similar results were obtained by our approach, with the major affiliation of the OTUs to the phyla *Firmicutes, Proteobacteria, and Bacteroidetes* (Supplementary Figure S14). We believe that our approach better succeeds to identify the core community for two reasons. First, our method identified core communities across assessments previously identified as not having a core community (as determined by 100% frequency occurrence). Second, our method offers a complement to other terms as “persistent’ and “transient” communities, e.g. indicating the rare microorganisms that could be classified in persistent group.

Same results were observed applying our proposed method to grapevine (leaves, flowers, grapes, and roots), and the maize rhizosphere. For example, Zarraonaindia *et al*., (2015) suggested a bacterial core community identified by three OTUs across 75% of samples from grape (leaves, flowers, grapes, and roots) and soils, over two growing seasons. These OTUs belonged to the genera *Bradyrhizobium, Steroidobacter* and *Acidobacteria*. By using our proposed method on the same dataset, 5,039 OTUs were identified as belonging to the core community (Supplementary Figure S12A and S12B). In addition, members of the *Cyanobacteria* phylum - which was a dominant group identified by the arbitrary methods (90% of relative abundance; Supplementary Figure S15) – comprised only a small component of the core microbial community using the probabilistic method. This variation in dominance could directly affect the conclusions about microbial composition across the system and may also affect the correlations with environmental drivers.

Here, we demonstrate the use of a probabilistic model to identify the core microbial communities. By applying a probabilistic model, our results suggest that the core microbial community may be higher in richness and diversity than previously demonstrated using other methods. Our method also allowed us to include rare (low abundance) members in the core microbial community, which would otherwise be a challenge using an arbitrary core cutoff. The use of a probabilistic model can extend our detection of the core microbial community, and could potentially help researchers to better connect the core community to ecosystem functions. An increased understanding of core microbial functions could support more robust studies in several fields, from human health (Zaura *et al*., 2009) to increased crop production. The microbial core community could also be used as an indicator of system perturbations (Shade and Handelsman, 2012) such as disease occurrence. This new approach could provide future studies a more realistic strategy to define calculate the core community, and could help to investigate the role of core microbial community in ecosystem function, or to elucidate the drivers of its composition. The probabilistic model is a new tool to step forward in the microbial community investigation. Only with the use of more rigorous and less arbitrary statistical methods it will be possible to understand the microbial ecology and its interactions.

## EXPERIMENTAL PROCEDURES

We selected four datasets composed of microbiomes from human samples (tongue, gut, and palms), mice (gut), grapevines (plant organs and bulk soil), and the maize rhizosphere to study the core microbial community identified using arbitrary cutoffs and a probabilistic method based on the Poisson distribution (Table 1).

**Table 1.**
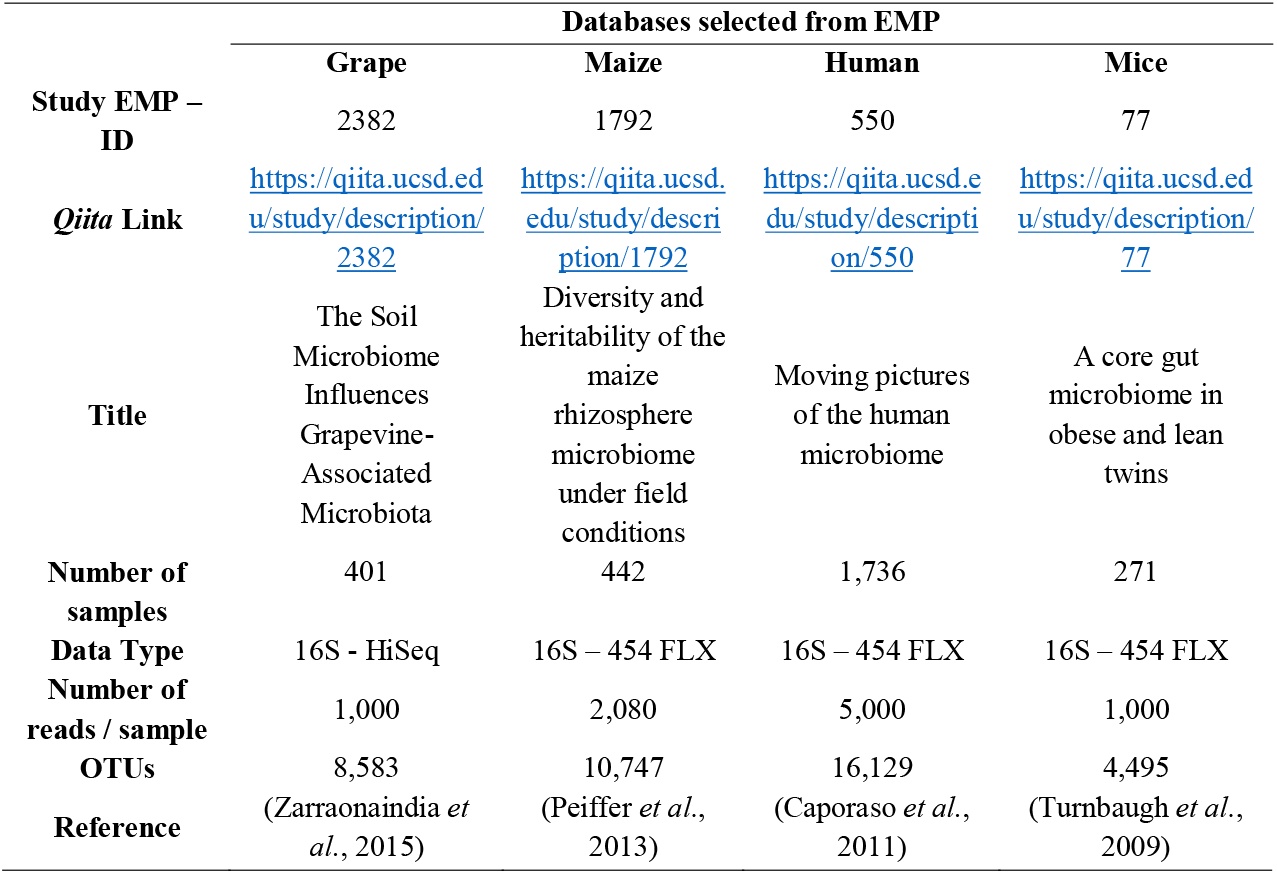
Databases selected from EMP on the *Qiita platform*.

**Table 2.** Number of OTU’s identified by the arbitrary and proposed method (based on the Poisson distribution) across the datasets

|  |  | Databases |  |  |  |  |  |  |  |
| --- | --- | --- | --- | --- | --- | --- | --- | --- | --- |
|  |  | Grapevine |  | Maize |  | Human |  | Mice |  |
| Methods |  | Core community | Variable community | Core community | Variable community | Core community | Variable community | Core community | Variable community |
| Conventional method | 30% | 211 | 8,372 | 272 | 10,475 | 206 | 15,923 | 170 | 4,325 |
|  | 40% | 109 | 8,474 | 145 | 10,602 | 93 | 16,036 | 82 | 4,413 |
|  | 50% | 40 | 8,543 | 80 | 10,667 | 42 | 16,087 | 35 | 4,460 |
|  | 60% | 15 | 8,568 | 39 | 10,708 | 24 | 16,105 | 19 | 4,476 |
|  | 70% | 5 | 8,578 | 19 | 10,728 | 12 | 16,117 | 5 | 4,490 |
|  | 80% | 0 | 8,583 | 5 | 10,742 | 2 | 16,127 | 2 | 4,493 |
|  | 90% | 0 | 8,583 | 3 | 10,744 | 0 | 16,129 | 0 | 4,495 |
|  | 100% | 0 | 8,583 | 0 | 10,747 | 0 | 16,129 | 0 | 4,495 |
| Proposed method |  | 5,039 | 3,544 | 5,294 | 5,453 | 8,751 | 7,378 | 1,717 | 2,778 |

The mice dataset was used to evaluate how the gut microbiome influences host adiposity (Turnbaugh *et al*., 2009). The data are from fecal samples from 154 individuals (mice) divided into adult females, monozygotic or dizygotic twin pairs, and their mothers. The core microbial community was identified using the *Phylotype Sampling Model* (PSM), which by Poisson distribution estimates the failures to observe microbial groups possibly belonging to the core community. The authors established a threshold value for abundance, considering only the OTUs with more than 0,5% of relative abundance as members of the core microbial community.

The human microbiome database consists of 396 samples, collected along a time series of two individuals at four body sites, including gut, tongue, and left and right palm (Caporaso *et al*., 2011). In the original study, the authors aimed to evaluate the temporal variation in the human microbiome. The authors used the terms persistent (microbial taxa with high levels of occurrence across samples), and transient (taxa with low levels of occurrence across samples) community, because it identified a very small temporal core across all samples. The core was defined as the taxa found across 100% of the samples.

In the grapevine database, Zarraonaindia *et al*., (2015) identify the OTUs shared across grapevine organs (flower, leaves, grapes, root), the root zone, and bulk soil over two growing seasons. The authors reduced the cutoff to 75% occurrence across samples to determine the core community. This decision was justified by the authors due to the lack of OTUs occurring across all samples.

The maize database is the only study included in our dataset that did not attempt to identify the core community. The authors aimed to determine the impact of genetic variation on the composition of bacterial communities inhabiting the maize rhizosphere (Peiffer *et al*., 2013).

The biological observation matrices (BIOM) derived from these data were obtained from the Earth Microbiome Project (EMP; http://www.earthmicrobiome.org), available on the *Qiita* platform (https://qiita.ucsd.edu). We used the BIOM files due to the similar treatment of data by bioinformatics, including quality filters and assignment of OTU taxonomy (Elli *et al*., 2010; Caporaso *et al*., 2011; Peiffer *et al*., 2013; Zarraonaindia *et al*., 2015). We used the software *QIIME* (Chen and Lifschitz, 1989) to convert the BIOM files into text files, which were further imported into the *R* software (Team 2016), where we analyzed it using the packages *‘RAM’* (Chen *et al*., 2016), *‘vegan’* (Oksanen *et al*., 2016) and *‘Hmisc’* (Harrell Jr *et al*., 2016).

The identification of the core microbial community is conventionally obtained by defining limits of frequency across the samples, *i.e*. a core community could be defined as microorganisms occurring in all samples (100% of occurrence frequency) or in a part of the samples (varying from 30% to 90% of frequency). For example, Ainsworth *et al*., (2015) identified the ubiquitous endosymbiont bacterial community (or core community) associated with corals using a 30% occurrence frequency cut-off. Similarly, the human and grapevine studies were used determined the core community, respectively at levels of 100%, 100% and 75% occurrence frequency across the samples. We used a range of limits - 30, 40, 50, 60,70, 80, 90 and 100% occurrence frequency - based on the OTU tables across the samples to verify the difference in the core microbial community selected by these methods.

The method proposed here is based on the probability test for the distribution of each microbial taxon (OTU) among samples. This probability test is based on the Poisson distribution, which is a discrete random probability regression model. The Poisson distribution expresses the probability of an event taking place at a given point in time (Rao and Rubin, 1964). Here we treat events as OTUs across a series of collected samples. The Poisson distribution has previously been used in biogeographic studies to predict the abundance of species in a given ecosystem (Vincent and Haworth, 1983; Guisan and Zimmermann, 2000).

Following the idea proposed in the Phylotype Sampling Model (Turnbaugh *et al*., 2009), the Poisson distribution was used to verify the sampling error expected given the sample size and the probability of observing the minimum abundance of a microorganism in any sample. However, the major difference from the previously methods including the Phylotype Sampling Model is that our proposed method does not present abundance or frequency thresholds. The probability *(**P**)* of Poisson distribution is obtained by ***P***(***x***) = ***λ^x^ e^−λ^***/***x!***, where the lambda (**λ**) and **x** represent the average of relative abundance and the occurrence frequency of each taxon across the communities, respectively. Using this formula, we have tested two hypotheses: ***H_0_*** – the individual (OTU) fits in the Poisson distribution and thus likely occurs in every sample (95% of confidence), indicating that it cannot be excluded from the core microbial community; ***H_1_*** – the individual does not fit in the Poisson distribution, and thus is unlikely to occur in every sample, supporting its exclusion from the core microbial community.

The calculation starts with the determination of the average of sequences per community source (***N***), the average relative abundance of each taxon across communities (***p***) and the occurrence frequency of each taxon across communities (**f**). The ***p*** and ***f*** are calculated with values of ***A*** and ***rich*** > 0, and they are used in the Poisson distribution, where the ***λ*** is obtained per OTU by the formula ***λ*** = ***N*** × ***p***.

The goodness-of-fit of the Poisson model to distribution of OTUs were determined from the R^2^ (adjusted) and *p-value*. The goodness of fit (R^2^) indicates the level of variance of an OTU’s relative abundance explained by the Poisson distribution, which in this case is correlated with the proportion of microbial community that could be not excluded as possible member of the core microbial group. The *p-value* is used to calculate the significance of OTUs predicted as probable core members by the Poisson distribution.

The arbitrary (thresholds of 30, 40, 50, 60, 70, 80, 90 and 100%) and the proposed (Poisson distribution) methods resulted in OTU tables for the core microbial community and the “variable” community (made of those that do not belong to the core community). The statistical analyses comparing the results were performed using the R software version 3.2.2 (R Core Team, 2015), including the Shannon index. We also developed a function in R, which identifies a core microbial community by the method based on the Poisson distribution. The R script of this function is available in Supplementary Code Simplified file, and the description is available in Supplementary Code Description file.

## Supporting information

## ACKNOWLEDGEMENTS

We thank FAPESP for the projects funding, 2014/22845-5 and 2013/18529-8. We are grateful to the students in Brendan J.M. Bohannan’s laboratory and the students from the Institute of Ecology and Evolution for support. We thank the members of the Earth Microbiome Project and the authors that made the database available. We also acknowledge the comments and discussion provided by Annelise Mendes Nascimento, Clarisse Betancourt, Ademir Durrer, and Trish Pasby throughout the manuscript preparation.

## Notes

***Conflict of Interest statement:*** The authors declare no conflict of interest.

