## Supplementary material for "A probabilistic model to identify the core microbial community"

Supplementary Code Description


### Supplementary Code Description

###### *Thiago Gumiere*

#### *2017-10-06*

### A probabilistic model to identify the core microbial community

The script named Supplementary\_Code\_simplified contains the script developed to identify the core microbial community and also export the graphic of the Poisson method. That script does not contain the conventional method identification.

This detail file contains the complete script (Poisson and conventional method). We will demonstrate using the four datasets the identification of the core microbial community. We also will build the graphics of Poisson and conventional methods.

#### Download the biom files from the Qiita platform

Download the biom files:

> Mice (https://qiita.ucsd.edu/study/description/77)

> Grape (https://qiita.ucsd.edu/study/description/2382)

> Human (https://qiita.ucsd.edu/study/description/550)

> Maize (https://qiita.ucsd.edu/study/description/1792)

After download and unzip the files from Qiita platform, the biom files should be rarefied in the same level indicated by each referent article (see Table 1). Here, we used the Qiime (http://qiime.org/) to rarefy the data sets. Using Qiime, the script used to rarefy sample is “single\_rarefaction.py” (http://qiime.org/scripts/single\_rarefaction.html). When the rarefaction is complete, the files can be converted to text format using “biom convert” (http://biom-format.org/documentation/biom\_conversion.html). This step will facilitate the import files in R.

#### Installing the packages required

The packages used as “Hmisc” (used during the Poisson model), “vegan” (multivariate analyses), and “RAM” (import .txt files).

```
#install.packages(c("Hmisc","vegan","RAM")) #It will take few minutes

require("Hmisc")
```

```
## Loading required package: Hmisc
```

```
## Loading required package: lattice
```

```
## Loading required package: survival
```

```
## Loading required package: Formula
```

```
## Loading required package: ggplot2
```

```
## 
## Attaching package: 'Hmisc'
```

```
## The following objects are masked from 'package:base':
## 
##     format.pval, units
```

```
require("vegan")
```

```
## Loading required package: vegan
```

```
## Loading required package: permute
```

```
## This is vegan 2.4-5
```

```
require("RAM")
```

```
## Loading required package: RAM
```

#### Importing rarefied files

The rarefied files are imported by “RAM” package using the function “fread.OTU”.

```
#Grape
setwd("~/Dropbox/Artigos/1_Core_concept/data_Core/grape/")
grape=fread.OTU("grape_otu_table_1000.txt")
```

```
## Warning in data.table::fread(file, na.strings = c("", ".", "NA")): Starting
## data input on line 2 and discarding line 1 because it has too few or too
## many items to be column names or data: # Constructed from biom file
```

```
#Human
setwd("~/Dropbox/Artigos/1_Core_concept/data_Core/human/")
human=fread.OTU("Human_otu_table_5000.txt")
```

```
## Warning in data.table::fread(file, na.strings = c("", ".", "NA")): Starting
## data input on line 2 and discarding line 1 because it has too few or too
## many items to be column names or data: # Constructed from biom file
```

```
## 
Read 62.0% of 16129 rows
Read 16129 rows and 1737 (of 1737) columns from 0.106 GB file in 00:00:04
```

```
#Maize
setwd("~/Dropbox/Artigos/1_Core_concept/data_Core/maize/")
maize=fread.OTU("maize_otu_table_2080.txt")
```

```
## Warning in data.table::fread(file, na.strings = c("", ".", "NA")): Starting
## data input on line 2 and discarding line 1 because it has too few or too
## many items to be column names or data: # Constructed from biom file
```

```
#MICE
setwd("~/Dropbox/Artigos/1_Core_concept/data_Core/mice/")
mice=fread.OTU("mice_otu_table_1000.txt")
```

```
## Warning in data.table::fread(file, na.strings = c("", ".", "NA")): Starting
## data input on line 2 and discarding line 1 because it has too few or too
## many items to be column names or data: # Constructed from biom file
```

```
#Resultados
setwd("~/Dropbox/Artigos/1_Core_concept/Results")
```

#### Running the Poisson model and identifying the core microbial community

We would like to emphazise that the Supplementary\_Code\_simplified file contains only the identification of core microbial community by the Poisson model (e.g. the Figure 1A). The script described below also contains the identification of the core by conventional methods (e.g. Figure 1B).

##### Script - Comparing Conventional with Poisson distribution

```
core.fit=function(otu_table) {
    otu_table_no_tax=otu_table[-length(otu_table)]
    otu_table_no_tax=t(otu_table_no_tax)
##Calculate the average of total individuals per community source

    N <- mean(apply(otu_table_no_tax, 1, sum))

##Calculate the average relative abundance of each taxa across communities
    p.m <- apply(otu_table_no_tax, 2, mean)
        p.m <- p.m[p.m != 0]
        p <- p.m/N

##Calculate the occurrence frequency of each taxa across communities
    freq <- apply(otu_table_no_tax>0, 2, mean)
    freq <- freq[freq != 0]
    
    
##Combine
    C <- merge(p, freq, by=0)
    C <- C[order(C[,2]),]
    C <- as.data.frame(C)
    C.0 <- C[!(apply(C, 1, function(y) any(y == 0))),] #Removes rows with any zero (absent in either source pool or local communities)
    p <- C.0[,2]
    freq <- C.0[,3]
    names(p) <- C.0[,1]
    names(freq) <- C.0[,1]

##Calculate the limit of detection  
    d = 1/N

##Goodness of fit for Poisson model
require(Hmisc)
    pois.pred <- ppois(d, N*p, lower.tail=F)
    pois.pred.ci<- binconf(pois.pred*nrow(otu_table), nrow(otu_table), alpha=0.95, method="exact", return.df=TRUE)
    
    otu_matrix=cbind(freq,p,pois.pred,pois.pred.ci)
    
    SS_res=sum((freq*log(freq/pois.pred))-(freq-pois.pred))
    SS_tot=sum((freq*log(freq/mean(freq))-(freq-mean(freq))))

    Rsqr.pois<- 1 - (SS_res/SS_tot)

# R square  adj
    SS_res2=sum((freq*log(freq/pois.pred))-(freq-pois.pred))+(length(subset(otu_matrix,freq>=pois.pred.ci[,1]))/2)
    
    SS_tot2=sum((freq*log(freq/mean(freq))-(freq-mean(freq))))


    Rsqr.pois.adj <- 1 - (SS_res2/SS_tot2)
    
# Root Mean Squared Error
    RMSE.pois =sum((pois.pred - freq)^2)/(length(freq) - 1)
    
#p-value
    t=(mean(freq)-pois.pred)/(sd(freq)/sqrt(length(freq)))
    p_value=2*pt(-abs(t),df=length(freq)-1)


## Split the the microbial core table and the variable community table
        core.table_Poisson=subset(otu_table,row.names(otu_table) %in% row.names(subset(otu_matrix,freq>=pois.pred.ci[,2])))
        core.table_30=subset(otu_table,row.names(otu_table) %in% row.names(subset(otu_matrix,freq>=0.3)))
        core.table_40=subset(otu_table,row.names(otu_table) %in% row.names(subset(otu_matrix,freq>=0.4)))
        core.table_50=subset(otu_table,row.names(otu_table) %in% row.names(subset(otu_matrix,freq>=0.5)))
        core.table_60=subset(otu_table,row.names(otu_table) %in% row.names(subset(otu_matrix,freq>=0.6)))
        core.table_70=subset(otu_table,row.names(otu_table) %in% row.names(subset(otu_matrix,freq>=0.7)))
        core.table_80=subset(otu_table,row.names(otu_table) %in% row.names(subset(otu_matrix,freq>=0.8)))
        core.table_90=subset(otu_table,row.names(otu_table) %in% row.names(subset(otu_matrix,freq>=0.9)))
        core.table_100=subset(otu_table,row.names(otu_table) %in% row.names(subset(otu_matrix,freq>=1.0)))
        
        
        nocore.table_Poisson=subset(otu_table,!row.names(otu_table) %in% row.names(subset(otu_matrix,freq>=pois.pred.ci[,2])))
        nocore.table_30=subset(otu_table,!row.names(otu_table) %in% row.names(subset(otu_matrix,freq>=0.3)))
        nocore.table_40=subset(otu_table,!row.names(otu_table) %in% row.names(subset(otu_matrix,freq>=0.4)))
        nocore.table_50=subset(otu_table,!row.names(otu_table) %in% row.names(subset(otu_matrix,freq>=0.5)))
        nocore.table_60=subset(otu_table,!row.names(otu_table) %in% row.names(subset(otu_matrix,freq>=0.6)))
        nocore.table_70=subset(otu_table,!row.names(otu_table) %in% row.names(subset(otu_matrix,freq>=0.7)))
        nocore.table_80=subset(otu_table,!row.names(otu_table) %in% row.names(subset(otu_matrix,freq>=0.8)))
        nocore.table_90=subset(otu_table,!row.names(otu_table) %in% row.names(subset(otu_matrix,freq>=0.9)))
        nocore.table_100=subset(otu_table,!row.names(otu_table) %in% row.names(subset(otu_matrix,freq>=1.0)))
        
return(list(core_Poisson=core.table_Poisson,core_30=core.table_30,core_40=core.table_40,core_50=core.table_50,core_60=core.table_60,core_70=core.table_70,core_80=core.table_80,core_90=core.table_90,core_100=core.table_100,nocore_Poisson=nocore.table_Poisson,nocore_30=nocore.table_30,nocore_40=nocore.table_40,nocore_50=nocore.table_50,nocore_60=nocore.table_60,nocore_70=nocore.table_70,nocore_80=nocore.table_80,nocore_90=nocore.table_90,nocore_100=nocore.table_100,otu_matrix=otu_matrix,Rsqr.pois.adj=Rsqr.pois.adj,p_value=p_value))
}
```

Running the script with each data set.

```
#Grape
grape_cores=core.fit(grape)

#Human
human_cores=core.fit(human)

#Maize
maize_cores=core.fit(maize)

#Mice
mice_cores=core.fit(mice)
```

The object “core\_Poisson” and “nocore\_Poisson” contains respectively the OTU table of core and variable microbial community.

For example, to isolate the core microbial community OTU table of mice dataset: **core.mice=mice\_cores$core\_Poisson**

For example, after the core identification the Figures 1, S10, S11, and S12 could be created.

#### Mice dataset

```
#Core figure
plot(freq~log(p),pch=18,col=ifelse(freq>=pois.pred,"blue","gray"), ylab="Occurence frequency",xlab="log(Mean Relative Abundance)",las=1,ylim=c(0,1),data=mice_cores$otu_matrix)

lines(pois.pred~log(p),col="red",lwd=2,mice_cores$otu_matrix) #Adding Poisson Line

text(-11.2,0.2,"core\ncommunity",col="blue")
text(-3.7,0.2,"variable\ncommunity",col="gray40")
text(-10,0.6, "Poisson distribution",font=2)
text(-10,0.5, bquote(bold(adj.R^2== .(round(mice_cores$Rsqr.pois.adj,digits=2)))))
text(-10,0.4, bquote(italic(p-value== .(round(mean(mice_cores$p_value),digits=3)))))
text(-12,1,"(A)")
```

*Figure 1A - The core and variable communities of the mice microbiome determined by (A) our proposed method based on the Poisson distribution.*

```
##Conventional figure
plot(freq~log(p),pch=18,col=ifelse(freq>=0.8,"blue",ifelse(freq>=0.7,"blue",ifelse(freq>=0.6,"blue",ifelse(freq>=0.5,"blue",ifelse(freq>=0.4,"blue",ifelse(freq>=0.3,"blue","gray")))))),ylab="Occurence frequency",las=1,xlab="log(Mean Relative Abundance)",mice_cores$otu_matrix,ylim=c(0,1))

lines(y=c(1,1),x=c(-9,0),lty=2,col="gray")
lines(y=c(0.9,0.9),x=c(-9,0),lty=2,col="gray")
lines(y=c(0.8,0.8),x=c(-9,0),lty=2,col="gray")
lines(y=c(0.7,0.7),x=c(-9,0),lty=2,col="gray")
lines(y=c(0.6,0.6),x=c(-9,0),lty=2,col="gray")
lines(y=c(0.5,0.5),x=c(-9,0),lty=2,col="gray")
lines(y=c(0.4,0.4),x=c(-9,0),lty=2,col="gray")
lines(y=c(0.3,0.3),x=c(-9,0),lty=2,col="gray")

text(-9,1.015,"100%",col="gray40",cex=0.8)
text(-9,0.915,"90%",col="gray40",cex=0.8)
text(-9,0.815,"80%",col="gray40",cex=0.8)
text(-9,0.715,"70%",col="gray40",cex=0.8)
text(-9,0.615,"60%",col="gray40",cex=0.8)
text(-9,0.515,"50%",col="gray40",cex=0.8)
text(-9,0.415,"40%",col="gray40",cex=0.8)
text(-9,0.315,"30%",col="gray40",cex=0.8)

text(-11.2,0.6,"core\ncommunity\n(thresholds)",col="blue")
text(-11.2,0.2,"variable\ncommunity",col="gray40")
text(-12,1,"(B)")
```

*Figure 1B – The core and variable communities of the mice microbiome determined by the conventional method.*

#### Human dataset

```
#Core figure
plot(freq~log(p),pch=18,col=ifelse(freq>=pois.pred,"blue","gray"), ylab="Occurence frequency",xlab="log(Mean Relative Abundance)",ylim=c(0,1),las=1,data=human_cores$otu_matrix)

lines(pois.pred~log(p),col="red",lwd=2,human_cores$otu_matrix) #Adding Poisson line

text(-14,0.15,"core\ncommunity",col="blue")
text(-4.5,0.10,"variable\ncommunity",col="gray40")
text(-13,0.6, "Poisson distribution",font=2,cex=0.9)
text(-13,0.5, bquote(bold(adj.R^2== .(round(human_cores$Rsqr.pois.adj,digits=2)))))
text(-13,0.4, bquote(italic(p-value== .(round(mean(human_cores$p_value),digits=3)))))
text(-15.9,1,"(A)")
```

*Figure S10A – The core and variable communities of the human microbiome determined by our proposed method based on the Poisson distribution.*

```
#Conventional figure
plot(freq~log(p),pch=18,col=ifelse(freq>=0.8,"blue",ifelse(freq>=0.7,"blue",ifelse(freq>=0.6,"blue",ifelse(freq>=0.5,"blue",ifelse(freq>=0.4,"blue",ifelse(freq>=0.3,"blue","gray")))))),ylab="Occurence frequency",las=1,xlab="log(Mean Relative Abundance)",human_cores$otu_matrix,ylim=c(0,1))

lines(y=c(1,1),x=c(-11,0),lty=2,col="gray")
lines(y=c(0.9,0.9),x=c(-11,0),lty=2,col="gray")
lines(y=c(0.8,0.8),x=c(-11,0),lty=2,col="gray")
lines(y=c(0.7,0.7),x=c(-11,0),lty=2,col="gray")
lines(y=c(0.6,0.6),x=c(-11,0),lty=2,col="gray")
lines(y=c(0.5,0.5),x=c(-11,0),lty=2,col="gray")
lines(y=c(0.4,0.4),x=c(-11,0),lty=2,col="gray")
lines(y=c(0.3,0.3),x=c(-11,0),lty=2,col="gray")

text(-11,1.015,"100%",col="gray40",cex=0.8)
text(-11,0.915,"90%",col="gray40",cex=0.8)
text(-11,0.815,"80%",col="gray40",cex=0.8)
text(-11,0.715,"70%",col="gray40",cex=0.8)
text(-11,0.615,"60%",col="gray40",cex=0.8)
text(-11,0.515,"50%",col="gray40",cex=0.8)
text(-11,0.415,"40%",col="gray40",cex=0.8)
text(-11,0.315,"30%",col="gray40",cex=0.8)

text(-14,0.6,"core\ncommunity\n(thresholds)",col="blue")
text(-14,0.2,"variable\ncommunity",col="gray40")
text(-15.9,1,"(B)")
```

*Figure S10B – The core and variable communities of the human microbiome determined by the conventional method.*

#### Grape dataset

```
#Core figure
plot(freq~log(p),pch=18,col=ifelse(freq>=pois.pred,"blue","gray"), ylab="Occurence frequency",xlab="log(Mean Relative Abundance)",ylim=c(0,1),las=1,data=grape_cores$otu_matrix)

lines(pois.pred~log(p),col="red",lwd=2,grape_cores$otu_matrix) #Adding Poisson Line

text(-11,0.15,"core\ncommunity",col="blue")
text(-4,0.15,"variable\ncommunity",col="gray40")
text(-10.2,0.6, "Poisson distribution",font=2,cex=0.9)
text(-10.5,0.5, bquote(bold(adj.R^2== .(round(grape_cores$Rsqr.pois.adj,digits=2)))))
text(-10.5,0.4, bquote(italic(p-value== .(round(mean(grape_cores$p_value),digits=3)))))
text(-12,1,"(A)")
```

*Figure S11A – The core and variable communities of the grape microbiome determined by our proposed method based on the Poisson distribution.*

```
#Conventional figure
plot(freq~log(p),pch=18,col=ifelse(freq>=0.8,"blue",ifelse(freq>=0.7,"blue",ifelse(freq>=0.6,"blue",ifelse(freq>=0.5,"blue",ifelse(freq>=0.4,"blue",ifelse(freq>=0.3,"blue","gray")))))),ylab="Occurence frequency",las=1,xlab="log(Mean Relative Abundance)",grape_cores$otu_matrix,ylim=c(0,1))

lines(y=c(1,1),x=c(-8.5,0),lty=2,col="gray")
lines(y=c(0.9,0.9),x=c(-8.5,0),lty=2,col="gray")
lines(y=c(0.8,0.8),x=c(-8.5,0),lty=2,col="gray")
lines(y=c(0.7,0.7),x=c(-8.5,0),lty=2,col="gray")
lines(y=c(0.6,0.6),x=c(-8.5,0),lty=2,col="gray")
lines(y=c(0.5,0.5),x=c(-8.5,0),lty=2,col="gray")
lines(y=c(0.4,0.4),x=c(-8.5,0),lty=2,col="gray")
lines(y=c(0.3,0.3),x=c(-8.5,0),lty=2,col="gray")

text(-8.5,1.015,"100%",col="gray40",cex=0.8)
text(-8.5,0.915,"90%",col="gray40",cex=0.8)
text(-8.5,0.815,"80%",col="gray40",cex=0.8)
text(-8.5,0.715,"70%",col="gray40",cex=0.8)
text(-8.5,0.615,"60%",col="gray40",cex=0.8)
text(-8.5,0.515,"50%",col="gray40",cex=0.8)
text(-8.5,0.415,"40%",col="gray40",cex=0.8)
text(-8.5,0.315,"30%",col="gray40",cex=0.8)

text(-11.5,0.6,"core\ncommunity\n(thresholds)",col="blue")
text(-11.5,0.2,"variable\ncommunity",col="gray40")
text(-12,1,"(B)")
```

*Figure S11B – The core and variable communities of the grape microbiome determined by the conventional method.*

#### Maize

```
#Core figure
plot(freq~log(p),pch=18,col=ifelse(freq>=pois.pred,"blue","gray"), ylab="Occurence frequency",xlab="log(Mean Relative Abundance)",ylim=c(0,1),las=1,data=maize_cores$otu_matrix)

lines(pois.pred~log(p),col="red",lwd=2,maize_cores$otu_matrix)

text(-12,0.15,"core\ncommunity",col="blue")
text(-4,0.15,"variable\ncommunity",col="gray40")
text(-11,0.6, "Poisson distribution",font=2,cex=0.9)
text(-11.3,0.5, bquote(bold(adj.R^2== .(round(maize_cores$Rsqr.pois.adj,digits=2)))))
text(-11.3,0.4, bquote(italic(p-value== .(round(mean(maize_cores$p_value),digits=3)))))
text(-13.5,1,"(A)")
```

*Figure S12A – The core and variable communities of the maize microbiome determined by our proposed method based on the Poisson distribution.*

```
#Conventional figure
plot(freq~log(p),pch=18,col=ifelse(freq>=0.8,"blue",ifelse(freq>=0.7,"blue",ifelse(freq>=0.6,"blue",ifelse(freq>=0.5,"blue",ifelse(freq>=0.4,"blue",ifelse(freq>=0.3,"blue","gray")))))),ylab="Occurence frequency",las=1,xlab="log(Mean Relative Abundance)",maize_cores$otu_matrix,ylim=c(0,1))

lines(y=c(1,1),x=c(-9,0),lty=2,col="gray")
lines(y=c(0.9,0.9),x=c(-9,0),lty=2,col="gray")
lines(y=c(0.8,0.8),x=c(-9,0),lty=2,col="gray")
lines(y=c(0.7,0.7),x=c(-9,0),lty=2,col="gray")
lines(y=c(0.6,0.6),x=c(-9,0),lty=2,col="gray")
lines(y=c(0.5,0.5),x=c(-9,0),lty=2,col="gray")
lines(y=c(0.4,0.4),x=c(-9,0),lty=2,col="gray")
lines(y=c(0.3,0.3),x=c(-9,0),lty=2,col="gray")

text(-9,1.015,"100%",col="gray40",cex=0.8)
text(-9,0.915,"90%",col="gray40",cex=0.8)
text(-9,0.815,"80%",col="gray40",cex=0.8)
text(-9,0.715,"70%",col="gray40",cex=0.8)
text(-9,0.615,"60%",col="gray40",cex=0.8)
text(-9,0.515,"50%",col="gray40",cex=0.8)
text(-9,0.415,"40%",col="gray40",cex=0.8)
text(-9,0.315,"30%",col="gray40",cex=0.8)

text(-11.7,0.6,"core\ncommunity\n(thresholds)",col="blue")
text(-11.7,0.2,"variable\ncommunity",col="gray40")
text(-13.5,1,"(B)")
```

*Figure S12B – The core and variable communities of the maize microbiome determined by the conventional method.*
