## Supplementary material for "A probabilistic model to identify the core microbial community"

```
## Core.fit Script Simplified##
## Authour: Thiago Gumiere ##
## 2017-10-06 ##

## The script below was developed to identify the core microbial community by the Poisson distribution.##
## This script was simplified not testing the conventional method.##
## This function requires the 'Hmisc' package ##
## The OTU table can be load by the package 'RAM' by the function 'fread.OTU'##

## The name_graphic in the functions indicates the name of the graphic which will be exported.##
## The name_graphic should be introduced between "quotation marks".##


core.fit=function(otu_table, name_graphic) {
	otu_table_no_tax=otu_table[-length(otu_table)]
	otu_table_no_tax=t(otu_table_no_tax)

	N <- mean(apply(otu_table_no_tax, 1, sum)) ##Calculate the average of total individuals per community source

	p.m <- apply(otu_table_no_tax, 2, mean) ##Calculate the average relative abundance of each taxa across communities
		p.m <- p.m[p.m != 0]
		p <- p.m/N

	freq <- apply(otu_table_no_tax>0, 2, mean) ##Calculate the occurrence frequency of each taxa across communities
	freq <- freq[freq != 0]
	
	C <- merge(p, freq, by=0) ##Combining the relative abundance and frequency
	C <- C[order(C[,2]),]
	C <- as.data.frame(C)
	C.0 <- C[!(apply(C, 1, function(y) any(y == 0))),] 

	#Removes rows with any zero (absent in either source pool or local communities)
	p <- C.0[,2] 
	freq <- C.0[,3]
	names(p) <- C.0[,1]
	names(freq) <- C.0[,1]

  
	d = 1/N ##Calculate the limit of detection	

  #Goodness of fit for Poisson model
  require(Hmisc)
	pois.pred <- ppois(d, N*p, lower.tail=F)
	pois.pred.ci<- binconf(pois.pred*nrow(otu_table), nrow(otu_table), alpha=0.95, method="exact", return.df=TRUE)
	
	otu_matrix=cbind(freq,p,pois.pred,pois.pred.ci)
	
	SS_res=sum((freq*log(freq/pois.pred))-(freq-pois.pred))
	SS_tot=sum((freq*log(freq/mean(freq))-(freq-mean(freq))))

	Rsqr.pois<- 1 - (SS_res/SS_tot)

  ##R square	adj
	SS_res2=sum((freq*log(freq/pois.pred))-(freq-pois.pred))+(length(subset(otu_matrix,freq>=pois.pred.ci[,1]))/2)
	
	SS_tot2=sum((freq*log(freq/mean(freq))-(freq-mean(freq))))


	Rsqr.pois.adj <- 1 - (SS_res2/SS_tot2)
	
  ##Root Mean Squared Error
	RMSE.pois =sum((pois.pred - freq)^2)/(length(freq) - 1)
	
  ##p-value
	t=(mean(freq)-pois.pred)/(sd(freq)/sqrt(length(freq)))
	p_value=2*pt(-abs(t),df=length(freq)-1)


  # Plot the microbial core commnuity (blue) and the variable community (gray)

	if(Rsqr.pois<=0) {
		stop('Negative or zero R-squared - It is not possible to determine the core and variable community')
		} else {
		pdf(file=paste("core_",name_graphic,".pdf", sep = ""),width=8,height=6)
		plot(freq~log(p),pch=18,col=ifelse(freq>=pois.pred.ci[,2],"blue","gray"), ylab="Occurence frequency",las=1,xlab="log(Mean Relative Abundance)",ylim=c(0,1),xlim=c(-17,-2),otu_matrix)
		lines(pois.pred~log(p),col="red",lwd=2)
		lines(pois.pred.ci[,2]~log(p),col="darkred",lty=2,lwd=1.5)
		text(-15,0.1,"core",col="blue")
		text(-4,0.1,"variable community",col="gray40")
		text(-15,0.6, "Poisson distribution",font=2)
		text(-15,0.5, bquote(bold(R^2== .(round(Rsqr.pois,digits=4)))))
		text(-15,0.4, bquote(bold(adj.R^2== .(round(Rsqr.pois.adj,digits=4)))))
		text(-15,0.45, bquote(italic(p-value== .(round(mean(p_value),digits=4)))))
		dev.off()

## Split the the microbial core table and the variable community table
		core.v=subset(otu_matrix,freq>=pois.pred.ci[,2])
		core.names=row.names(core.v)

		index1=row.names(otu_table) %in% core.names
		core.table=subset(otu_table,index1)
	
		index2=!row.names(otu_table) %in% core.names
		nocore.table=subset(otu_table,index2)
		return(list(core.table=core.table,nocore.table=nocore.table,pois.pred=otu_matrix))
	}
}

## core.table contains the core microbial community identified by the poisson method
## nocore.table constains the variable community
## pois.pred contains frequency and abundance of OTU's, and the predicted values of Poisson distribution
```
