## Supplementary material for "A probabilistic model to identify the core microbial community"

***Running header****: Identifying the probable core microbial community*

***Authors: ^1*^****Thiago Gumiere, ^2^Kyle Meyer, ^2^Adam Burns, ^3^Silvio J. Gumiere, ^2^Brendan J. M. Bohannan, ^1^Fernando D. Andreote*

1. ***Testing models to select probable members of the core microbial community***

We tested 13 different distribution models (listed below) to verify if the Poisson fit better in datasets. The models were applied for the OTU’s from each database, available in *Qiita plataform*. Also, we have used the same rarefied levels and assign references indicated by each work published.

•	Beta distribution;

•	Cauchy distribution;

•	Chi-squared distribution;

•	Exponential distribution;

•	Gamma distribution;

•	Geometric distribution;

•	Log-normal distribution;

•	Negative binomial;

•	Normal distribution;

•	Poisson distribution;

•	Student's t distribution;

•	Uniform distribution;

•	Weibull distribution.

The selection was based on the fitted R^2^, Root-mean-square error (RMSE), and p-value. As observed in Table S1, the Poisson distribution showed the higher R^2^ and *p-value < 0.05*, and lesser value of RMSE. We selected the four distribution models with higher R^2^ and showed their fitting on OTU datasets.

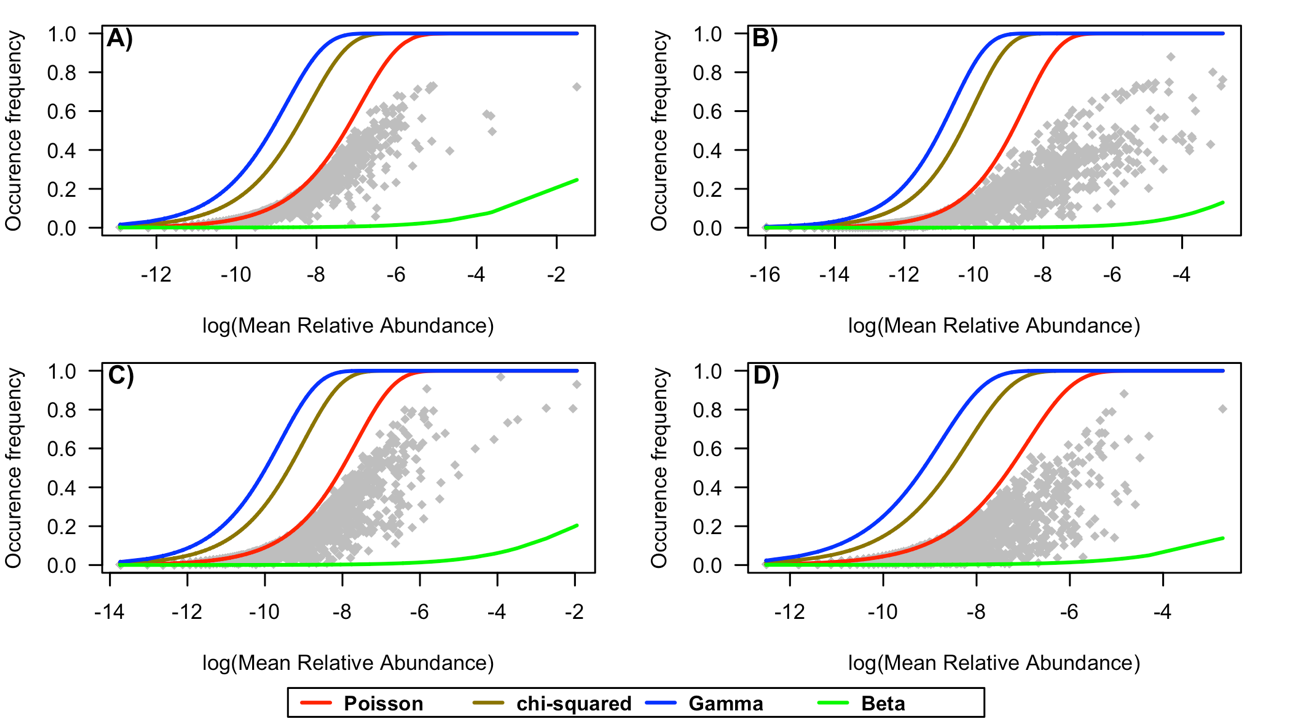

Figure S1 - Probabilistic models fitted on the OTU’s distribution. The Poisson, Chi-squared, Gamma and Beta distribution was the models with higher R^2^, lower RSME, and significant p-value (>0.05). The datasets Grape, Human, Mice and Maize are respectively indicated by the letters A, B, C and D.

Table S1 – Comparing different distribution model fit in OTU tables from analyzed datasets. The models are classified by R^2^, Root-mean-square error (RMSE), and significance of p-value.

|  | **Maize** | | | |  | | **Human** | | | |  | | **Grape** | | | |  | | **Mice** | | |
| --- | --- | --- | --- | --- | --- | --- | --- | --- | --- | --- | --- | --- | --- | --- | --- | --- | --- | --- | --- | --- | --- |
| **Distribution Models** | **R^2^** | **RMSE** | **p-value** |  | | **R^2^** | | **RMSE** | **p-value** |  | | **R^2^** | | **RMSE** | **p-value** |  | | **R^2^** | | **RMSE** | **p-value** |
| Poisson | 0.83763 | 0.00595 | <0.05 |  | | 0.69063 | | 0.01094 | <0.05 |  | | 0.91813 | | 0.00294 | <0.05 |  | | 0.4778 | | 0.01944 | <0.05 |
| Chi-squared | -0.50513 | 0.0473 | <0.05 |  | | -0.36832 | | 0.03654 | <0.05 |  | | -0.17622 | | 0.03197 | <0.05 |  | | -1.30541 | | 0.09097 | <0.05 |
| Gamma | -1.71562 | 0.08886 | <0.05 |  | | -1.22933 | | 0.05741 | <0.05 |  | | -1.33235 | | 0.067 | <0.05 |  | | -2.68746 | | 0.15161 | <0.05 |
| Beta | -2.28047 | 0.00925 | <0.05 |  | | -1.22372 | | 0.0043 | <0.05 |  | | -1.85309 | | 0.00818 | <0.05 |  | | -2.19261 | | 0.01249 | <0.05 |
| Negative Binomial | -5.78976 | 0.00955 | <0.05 |  | | -6.74792 | | 0.00451 | <0.05 |  | | -3.79877 | | 0.00846 | <0.05 |  | | -5.25607 | | 0.01312 | <0.05 |
| Weibull | -7.08573 | 0.21538 | <0.05 |  | | -9.65882 | | 0.18805 | <0.05 |  | | -7.26305 | | 0.19642 | <0.05 |  | | -6.74743 | | 0.26717 | <0.05 |
| Student’s t | -7.60175 | 0.21434 | <0.05 |  | | -11.57893 | | 0.23209 | <0.05 |  | | -8.15619 | | 0.21438 | <0.05 |  | | -6.18332 | | 0.20203 | <0.05 |
| Normal | -7.60247 | 0.21437 | <0.05 |  | | -11.57919 | | 0.2321 | <0.05 |  | | -8.15771 | | 0.21445 | <0.05 |  | | -6.18511 | | 0.20212 | <0.05 |
| Exponential | -17.89706 | 0.91911 | <0.05 |  | | -25.32996 | | 0.95964 | <0.05 |  | | -19.13756 | | 0.92024 | <0.05 |  | | -15.30236 | | 0.89081 | <0.05 |
| Geometric | -17.89706 | 0.91911 | <0.05 |  | | -25.32996 | | 0.95964 | <0.05 |  | | -19.13756 | | 0.92024 | <0.05 |  | | -15.30235 | | 0.89081 | <0.05 |
| Uniform | -17.8974 | 0.91914 | <0.05 |  | | -25.33072 | | 0.95969 | <0.05 |  | | -19.13794 | | 0.92028 | <0.05 |  | | -15.30382 | | 0.89095 | <0.05 |
| Cauchy | -17.89798 | 0.9192 | <0.05 |  | | -25.33089 | | 0.9597 | <0.05 |  | | -19.13914 | | 0.92038 | <0.05 |  | | -15.3054 | | 0.89111 | <0.05 |
| Log-normal | -17.89798 | 0.9192 | <0.05 |  | | -25.33089 | | 0.9597 | <0.05 |  | | -19.13915 | | 0.92038 | <0.05 |  | | -15.30541 | | 0.89111 | <0.05 |

2. **Testing the rarefaction effect on Poisson distribution model and core microbial community**

As the Poisson distribution requires an equal number of reads per sample, we tested the effect of rarefication method. The Figures S2, S3, S4, and S5 refers to the Poisson fitting under different rarefactions levels. The small graphic inside of each figure indicated the adjusted R^2^ of each Poisson line under different rarefactions levels. The next group of figures (Figures S6, S7, S8, and S9) indicates the taxonomic composition of the core microbial community resulted from each rarefication level. The Table S2 indicates the values of samples, observation (OTU’s), total count (total numbers of reads) of the original OTU table (obtained from Qiita plataform). The Table S2 also indicates the adjusted R^2^, RMSE, and number of OTU’s identified by the Poisson distribution. The Figure S10 indicates the NMDS analyses for each dataset comparing the core microbial community identified

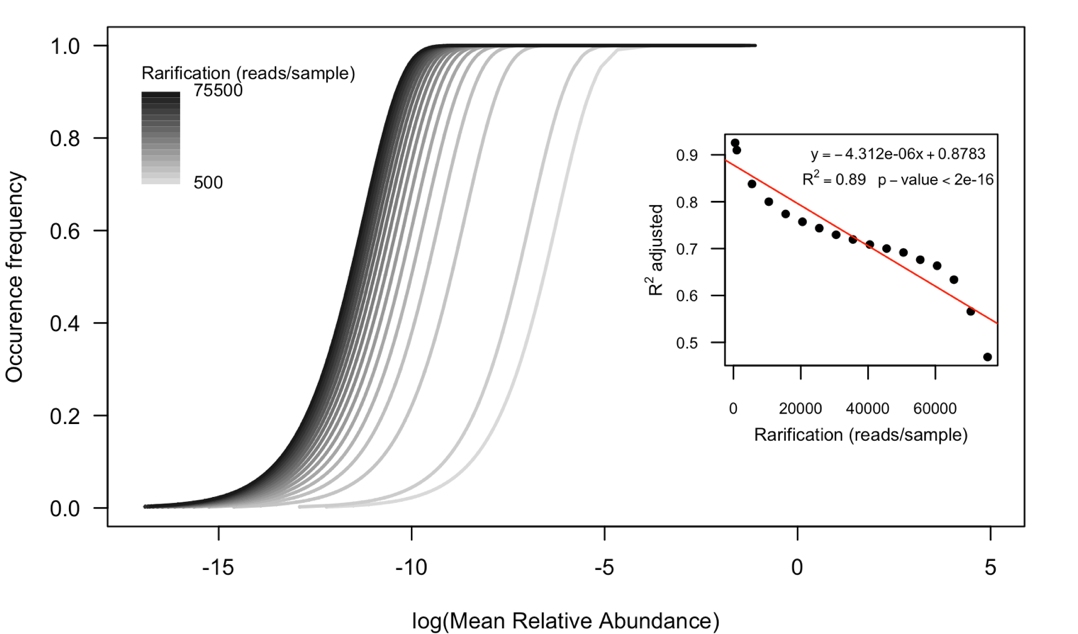

Figure S2 – Evaluating the rarefaction effect under the Poisson distribution fit. The grape dataset was rarefied from 500 to 75500 reads per sample. The grayscale lines indicated each Poisson line applied to specific rarefaction level. The small graphic inside indicates the tendency of adjusted R^2^ of Poisson line under different rarefication levels.

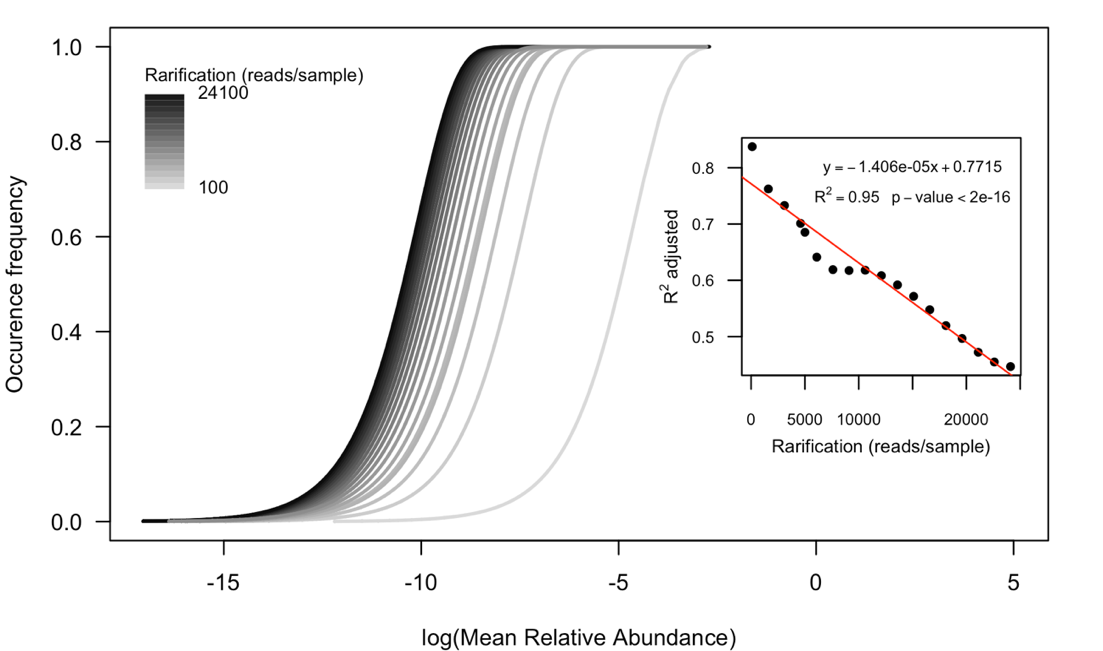

Figure S3 – Evaluating the rarefaction effect under the Poisson distribution fit. The human dataset was rarefied from 100 to 24100 reads per sample. The grayscale lines indicated each Poisson line applied to specific rarefaction level. The small graphic inside indicates the tendency of adjusted R^2^ of Poisson line under different rarefication levels.

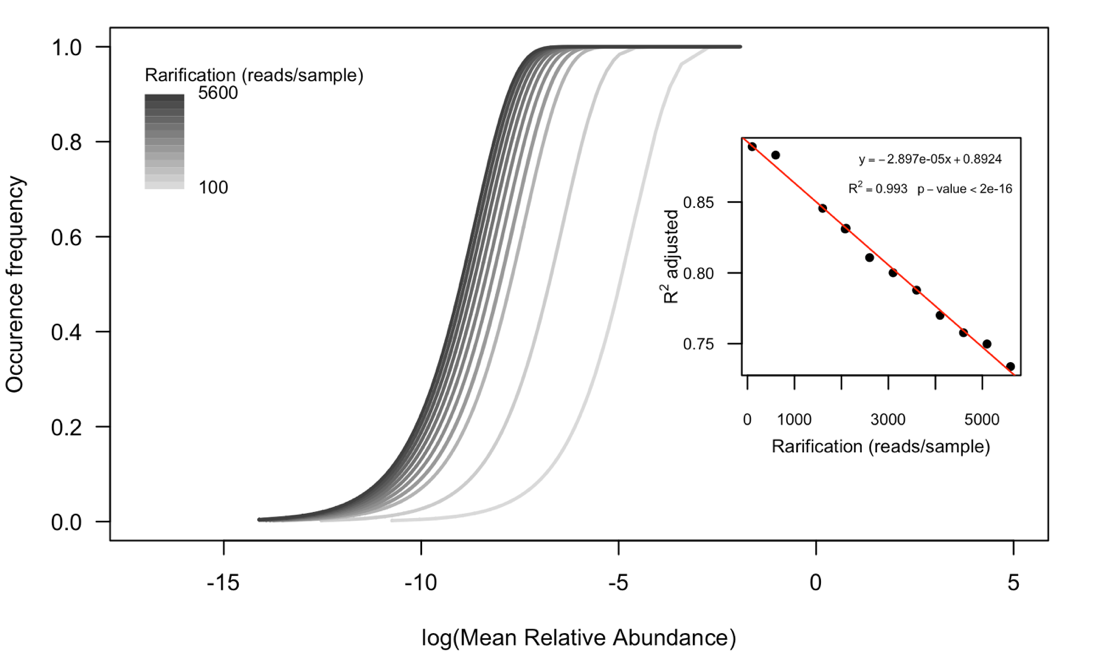

Figure S4 – Evaluating the rarefaction effect under the Poisson distribution fit. The maize dataset was rarefied from 100 to 5600 reads per sample. The grayscale lines indicated each Poisson line applied to specific rarefaction level. The small graphic inside indicates the tendency of adjusted R^2^ of Poisson line under different rarefication levels.

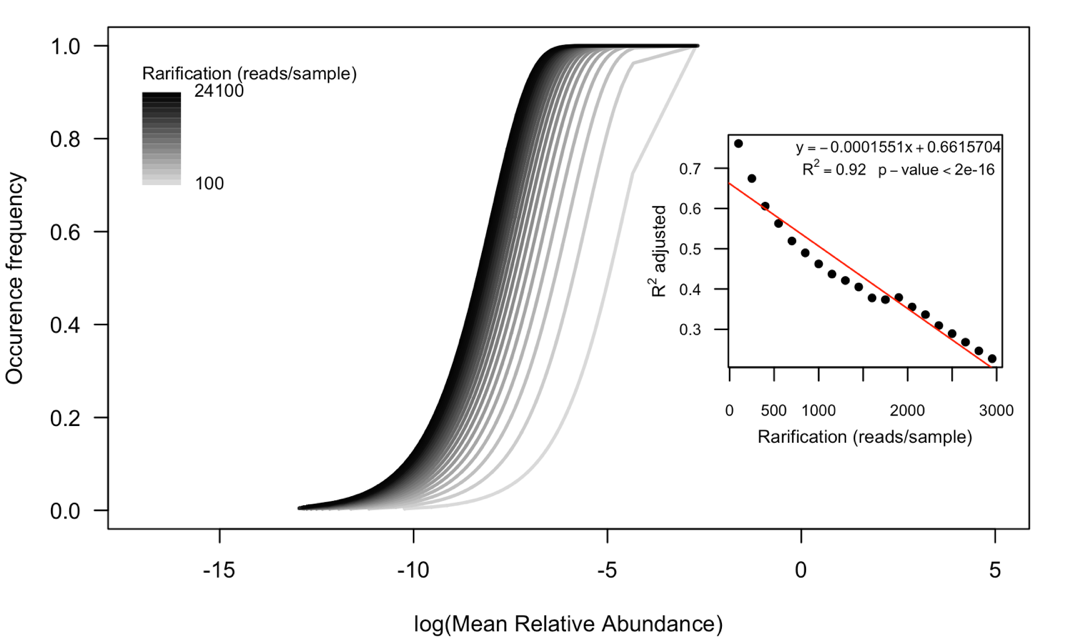

Figure S5 – Evaluating the rarefaction effect under the Poisson distribution fit. The mice dataset was rarefied from 100 to 24100 reads per sample. The grayscale lines indicated each Poisson line applied to specific rarefaction level. The small graphic inside indicates the tendency of adjusted R^2^ of Poisson line under different rarefication levels.

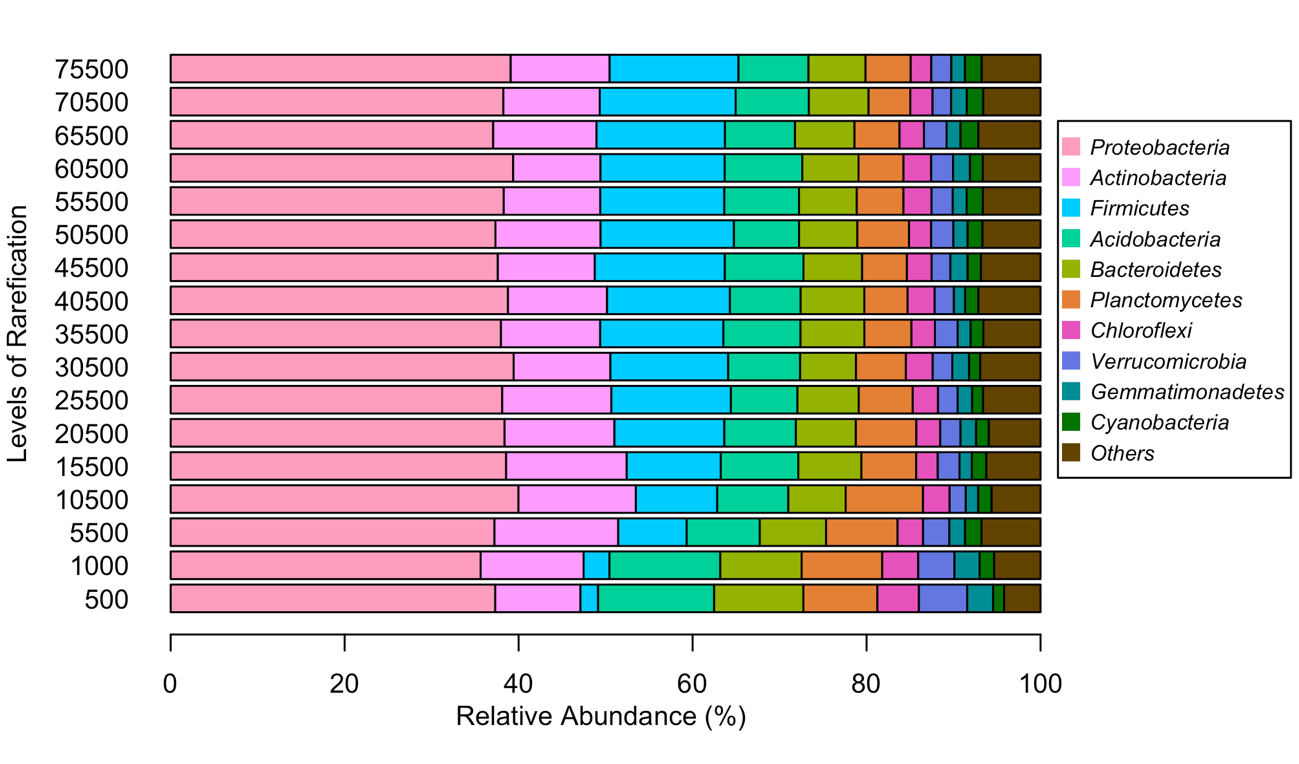

Figure S6 – Evaluating the rarefaction effect under the core microbial community identified by the Poisson distribution of grape dataset. The taxonomic composition is showed at the Phylum level.

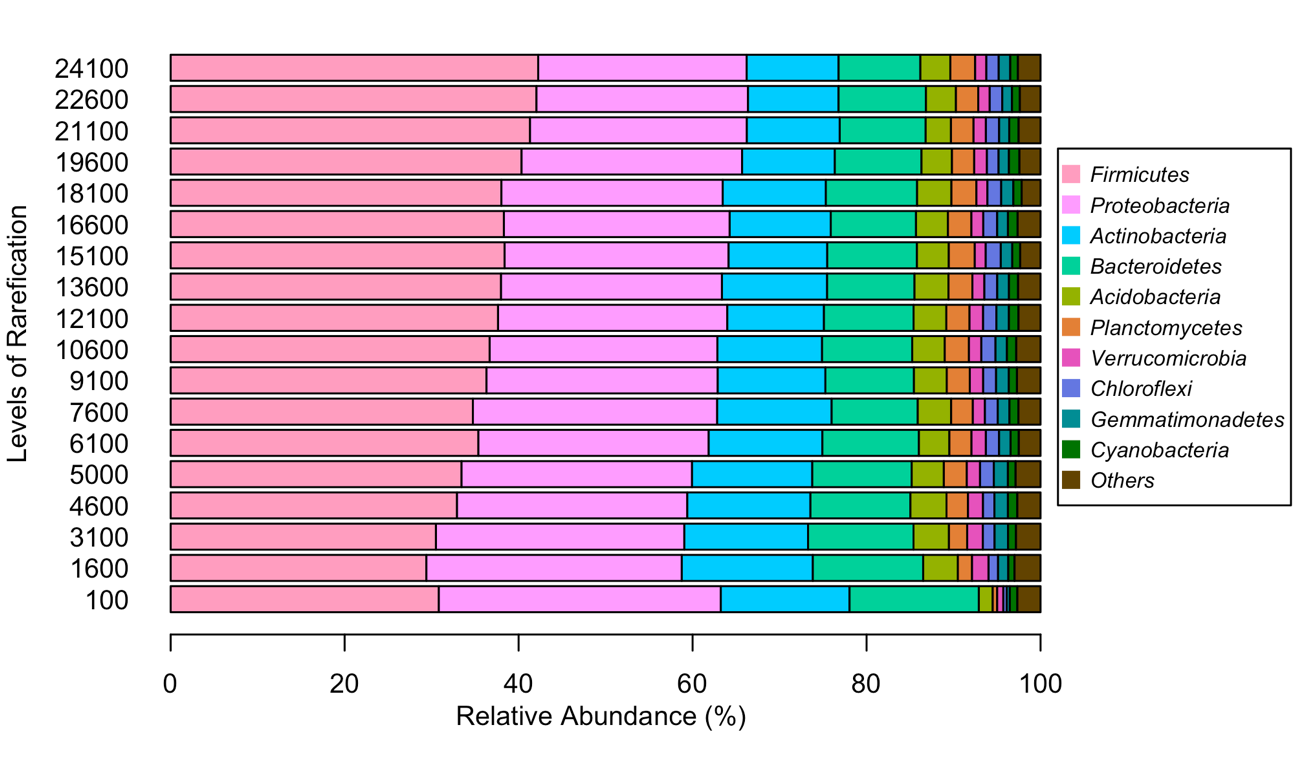

Figure S7 – Evaluating the rarefaction effect under the core microbial community identified by the Poisson distribution of human dataset. The taxonomic composition is showed at the Phylum level.

.

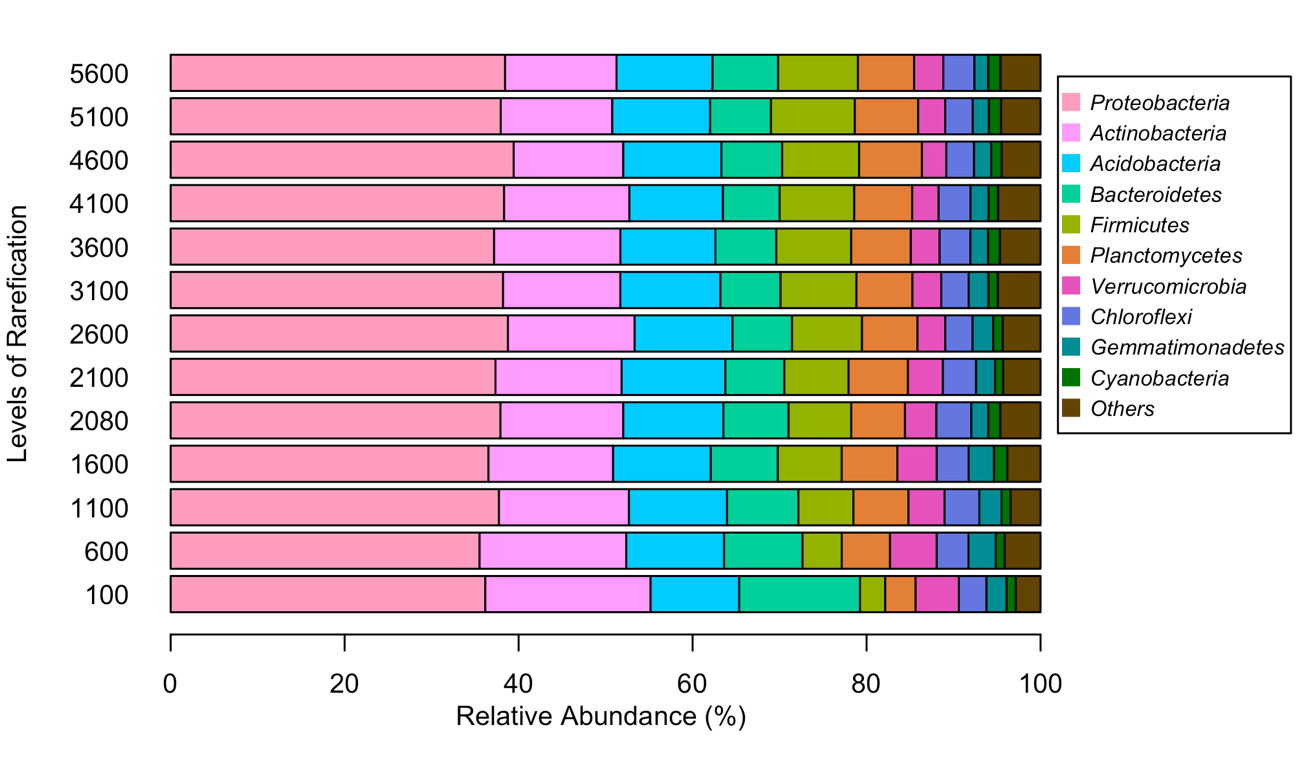

Figure S8 – Evaluating the rarefaction effect under the core microbial community identified by the Poisson distribution of maize dataset. The taxonomic composition is showed at the Phylum level

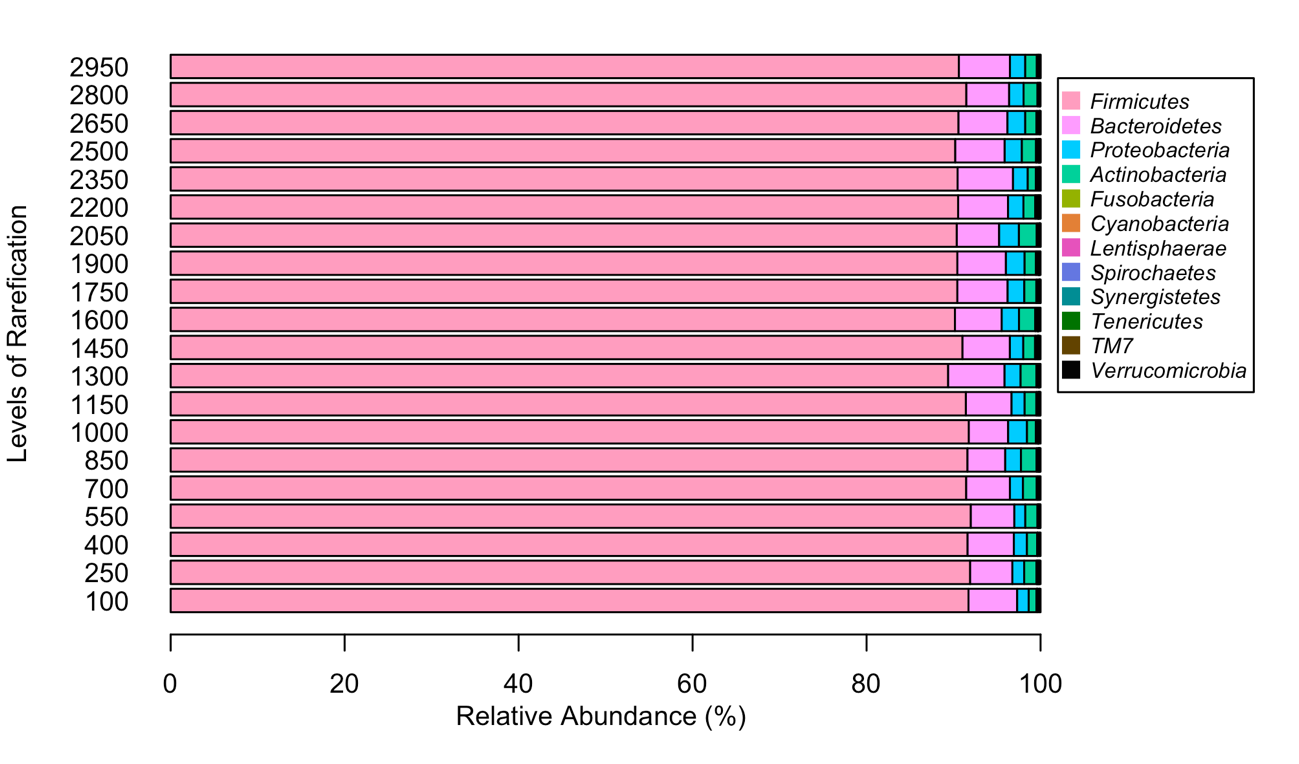

Figure S9 – Evaluating the rarefaction effect under the core microbial community identified by the Poisson distribution of mice dataset. The taxonomic composition is showed at the Phylum level

**Table S2 - Rarefaction effect on the original microbial community (original OTU table) and on the core microbial community identified by the Poisson distribution. The highlight values (italic and bold) are outliers with significant difference from the other values in their datasets (Chi-square, p-value<0.05).**

|  | **Rarefaction** | | | | | **Poisson Distribution** | | |
| --- | --- | --- | --- | --- | --- | --- | --- | --- |
|  | **Rarefaction level** | **Num. samples** | **Num. Observations** | **Total count** | **Table density*** | **R^2^** | **RMSE** | **OTU’s - Core** |
| **Grape** | 500 | 400 | 7195 | 200000 | 0.032 | 0.93 | 0.001 | 4895 |
|  | 1000 | 400 | 8583 | 400000 | 0.044 | 0.91 | 0.003 | 5039 |
|  | 5500 | 400 | 12055 | 2200000 | 0.088 | 0.84 | 0.013 | 5156 |
|  | 10500 | 400 | 13505 | 4200000 | 0.108 | 0.80 | 0.019 | 5081 |
|  | 15500 | 400 | 14285 | 6200000 | 0.121 | 0.77 | 0.025 | 5092 |
|  | 20500 | 400 | 14959 | 8200000 | 0.129 | 0.76 | 0.029 | 4993 |
|  | 25500 | 400 | 15483 | 10200000 | 0.136 | 0.74 | 0.032 | 5080 |
|  | 30500 | 398 | 15827 | 12139000 | 0.142 | 0.73 | 0.035 | 5126 |
|  | 35500 | 396 | 16199 | 14058000 | 0.146 | 0.72 | 0.037 | 5055 |
|  | 40500 | 395 | 16454 | 15997500 | 0.15 | 0.71 | 0.039 | 5029 |
|  | 45500 | 392 | 16754 | 17836000 | 0.153 | 0.70 | 0.041 | 5044 |
|  | 50500 | 388 | 16938 | 19594000 | 0.157 | 0.69 | 0.043 | 4968 |
|  | 55500 | 374 | 17132 | 20757000 | 0.159 | 0.68 | 0.046 | 4959 |
|  | 60500 | 364 | 17340 | 22022000 | 0.161 | 0.66 | 0.048 | 4974 |
|  | 65500 | 334 | 17233 | 21877000 | 0.163 | 0.63 | 0.052 | 4859 |
|  | ***70500*** | ***288*** | ***16927*** | ***20304000*** | ***0.16*** | ***0.57*** | ***0.058*** | ***4741*** |
|  | ***75500*** | ***249*** | ***16616*** | ***18799500*** | ***0.153*** | ***0.47*** | ***0.065*** | ***4530*** |
| **Maize** | ***100*** | ***460*** | ***4345*** | ***46000*** | ***0.012*** | ***0.89*** | ***0.001*** | ***3668*** |
|  | 600 | 459 | 8017 | 275400 | 0.026 | 0.88 | 0.002 | 5175 |
|  | 1100 | 459 | 9348 | 504900 | 0.034 | 0.86 | 0.003 | 5331 |
|  | 1600 | 458 | 10163 | 732800 | 0.04 | 0.85 | 0.005 | 5301 |
|  | 2080 | 441 | 10747 | 917280 | 0.045 | 0.83 | 0.006 | 5294 |
|  | 2100 | 441 | 10771 | 926100 | 0.045 | 0.83 | 0.006 | 5312 |
|  | 2600 | 389 | 10944 | 1011400 | 0.051 | 0.81 | 0.007 | 5188 |
|  | 3100 | 361 | 11203 | 1119100 | 0.055 | 0.80 | 0.008 | 5090 |
|  | 3600 | 330 | 11318 | 1188000 | 0.06 | 0.79 | 0.010 | 5044 |
|  | 4100 | 299 | 11451 | 1225900 | 0.064 | 0.77 | 0.011 | 5000 |
|  | 4600 | 280 | 11429 | 1288000 | 0.067 | 0.76 | 0.012 | 4899 |
|  | 5100 | 261 | 11503 | 1331100 | 0.07 | 0.75 | 0.013 | 4921 |
|  | 5600 | 238 | 11365 | 1332800 | 0.073 | 0.73 | 0.014 | 4764 |
| **Human** | ***100*** | ***1967*** | ***4828*** | ***196700*** | ***0.008*** | 0.84 | 0.002 | ***3866*** |
|  | 1600 | 1955 | 12846 | 3128000 | 0.017 | 0.76 | 0.006 | 8291 |
|  | 3100 | 1936 | 15206 | 6001600 | 0.02 | 0.73 | 0.008 | 8788 |
|  | 4600 | 1802 | 16052 | 8289200 | 0.022 | 0.70 | 0.010 | 8673 |
|  | 5000 | 1735 | 16129 | 8675000 | 0.022 | 0.69 | 0.011 | 8751 |
|  | 6100 | 1571 | 15993 | 9583100 | 0.023 | 0.64 | 0.013 | 8273 |
|  | 7600 | 1516 | 16539 | 11521600 | 0.025 | 0.62 | 0.014 | 8166 |
|  | 9100 | 1435 | 17270 | 13058500 | 0.026 | 0.62 | 0.015 | 8165 |
|  | 10600 | 1360 | 17874 | 14416000 | 0.028 | 0.62 | 0.015 | 8191 |
|  | 12100 | 1323 | 18214 | 16008300 | 0.03 | 0.61 | 0.016 | 7972 |
|  | 13600 | 1294 | 18610 | 17598400 | 0.03 | 0.59 | 0.017 | 8043 |
|  | 15100 | 1268 | 18801 | 19146800 | 0.031 | 0.57 | 0.018 | 7897 |
|  | 16600 | 1225 | 18691 | 20335000 | 0.031 | 0.55 | 0.019 | 7704 |
|  | 18100 | 1182 | 18405 | 21394200 | 0.031 | 0.52 | 0.020 | 7548 |
|  | 19600 | 1128 | 18018 | 22108800 | 0.032 | 0.50 | 0.021 | 7329 |
|  | 21100 | 1089 | 17803 | 22977900 | 0.031 | 0.47 | 0.022 | 7103 |
|  | 22600 | 1061 | 17727 | 23978600 | 0.032 | 0.45 | 0.023 | 6944 |
|  | 24100 | 1041 | 17771 | 25088100 | 0.032 | 0.45 | 0.023 | 6894 |
| **Mice** | 100 | 278 | 2655 | 27800 | 0.023 | 0.76 | 0.001 | 1699 |
|  | 250 | 278 | 3394 | 69500 | 0.035 | 0.67 | 0.004 | 1797 |
|  | 400 | 277 | 3782 | 110800 | 0.042 | 0.61 | 0.008 | 1815 |
|  | 550 | 276 | 4043 | 151800 | 0.048 | 0.56 | 0.011 | 1775 |
|  | 700 | 274 | 4229 | 191800 | 0.053 | 0.52 | 0.014 | 1717 |
|  | 850 | 273 | 4381 | 232050 | 0.057 | 0.49 | 0.017 | 1713 |
|  | 1000 | 270 | 4485 | 270000 | 0.061 | 0.46 | 0.020 | 1658 |
|  | 1150 | 270 | 4589 | 310500 | 0.065 | 0.44 | 0.022 | 1658 |
|  | 1300 | 267 | 4669 | 347100 | 0.068 | 0.42 | 0.025 | 1622 |
|  | 1450 | 257 | 4714 | 372650 | 0.071 | 0.41 | 0.027 | 1587 |
|  | 1600 | 244 | 4744 | 390400 | 0.074 | 0.38 | 0.029 | 1567 |
|  | 1750 | 233 | 4774 | 407750 | 0.077 | 0.37 | 0.031 | 1589 |
|  | 1900 | 222 | 4790 | 421800 | 0.081 | 0.38 | 0.033 | 1571 |
|  | 2050 | 200 | 4731 | 410000 | 0.085 | 0.36 | 0.036 | 1539 |
|  | 2200 | 189 | 4751 | 415800 | 0.088 | 0.34 | 0.038 | 1509 |
|  | 2350 | 167 | 4661 | 392450 | 0.093 | 0.31 | 0.040 | 1474 |
|  | 2500 | 153 | 4617 | 382500 | 0.096 | 0.29 | 0.043 | 1476 |
|  | 2650 | 140 | 4551 | 371000 | 0.1 | 0.27 | 0.045 | 1379 |
|  | 2800 | 123 | 4468 | 344400 | 0.105 | 0.25 | 0.048 | 1386 |
|  | 2950 | 112 | 4430 | 330400 | 0.109 | 0.23 | 0.051 | 1389 |

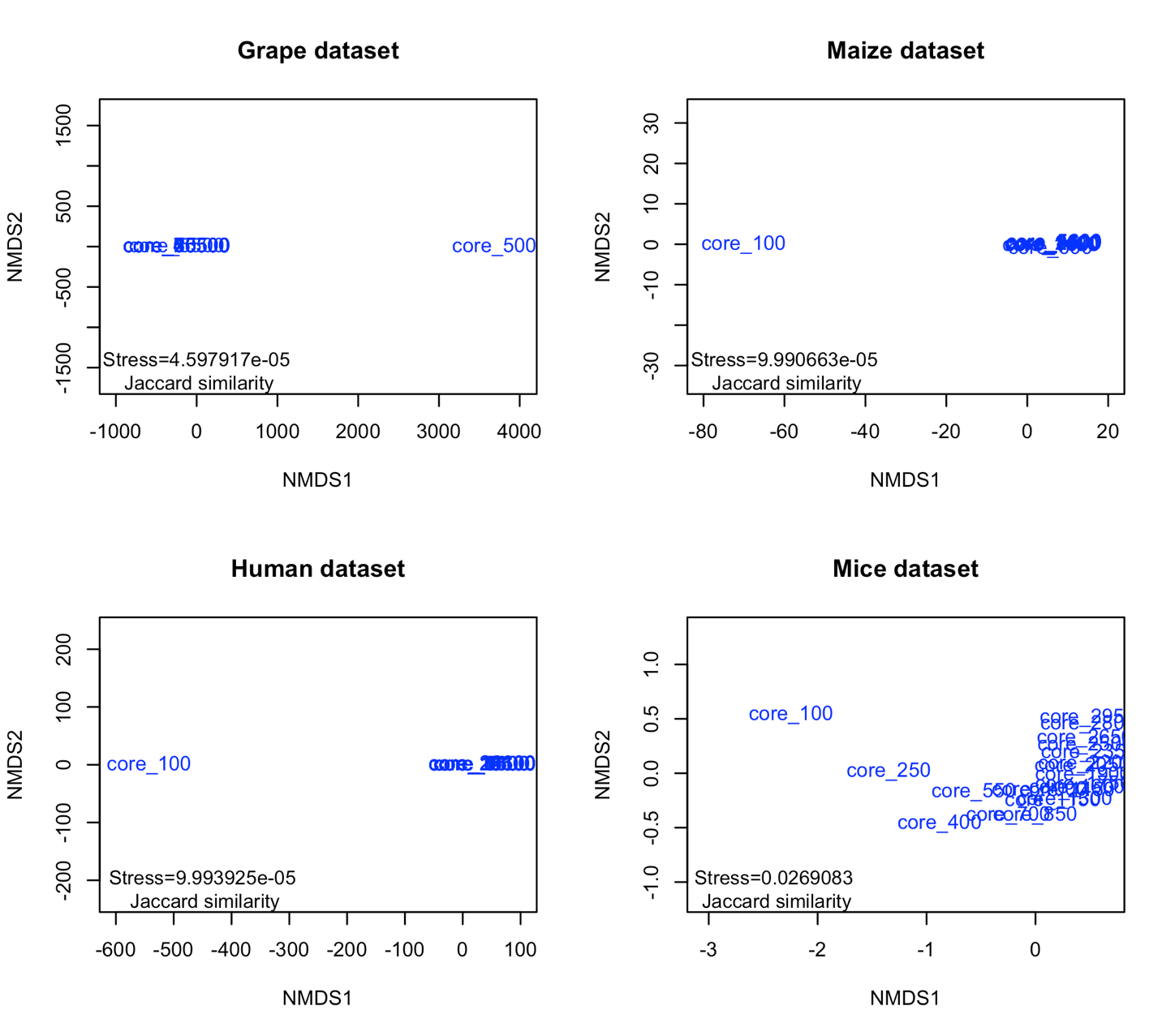

Figure S10 – NMDS of rarefaction effect on the core microbial community composition identified by the Poisson distribution for each dataset.

3. **Identifying the core microbial community by the Poisson distribution model and conventional method**

As indicated in Figures S11, S12 and S13, we compared the identification of core microbial community by the Poisson method (A) with the conventional method (B) for each dataset. The next group of figures (Figures S14, S15, and S16) are related with the taxonomic composition of each core microbial community identified by Poisson and conventional method.

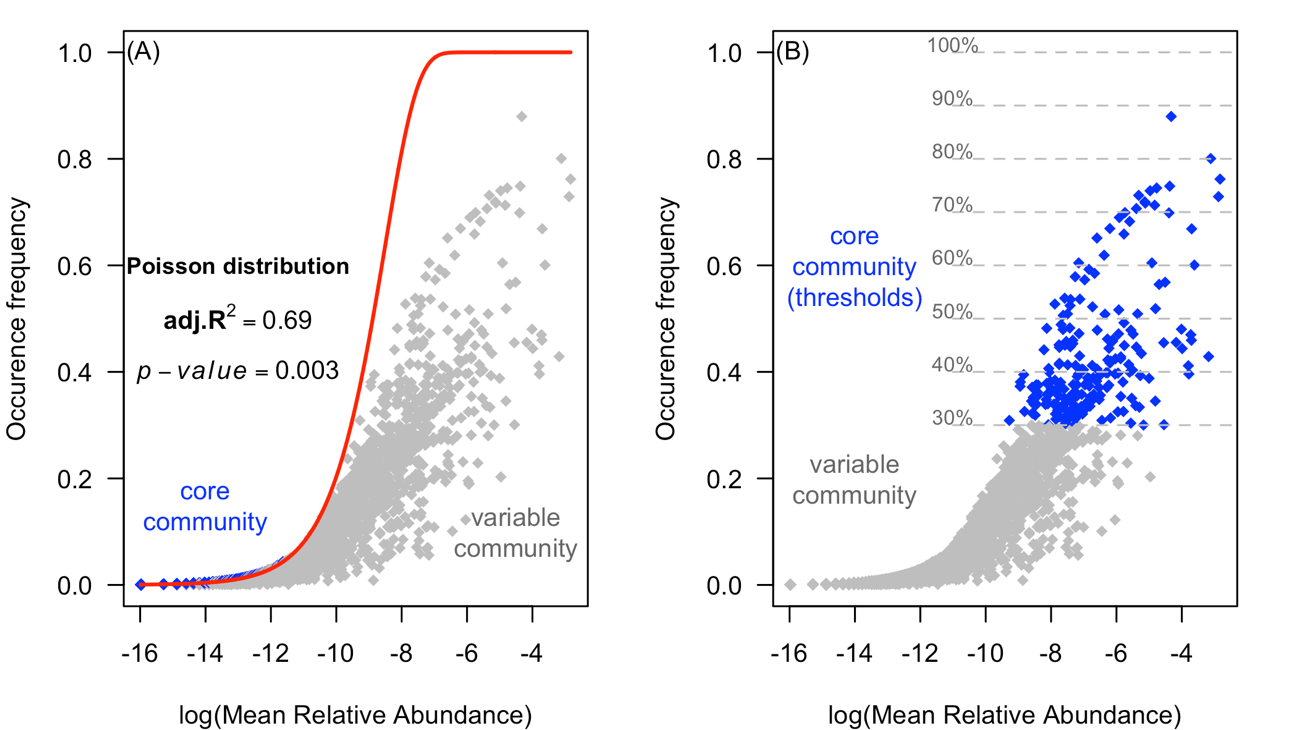

Figure S11 – The core and variable communities of the human microbiome determined by (A) our proposed method based on the Poisson distribution and (B) an arbitrary, threshold-based method.

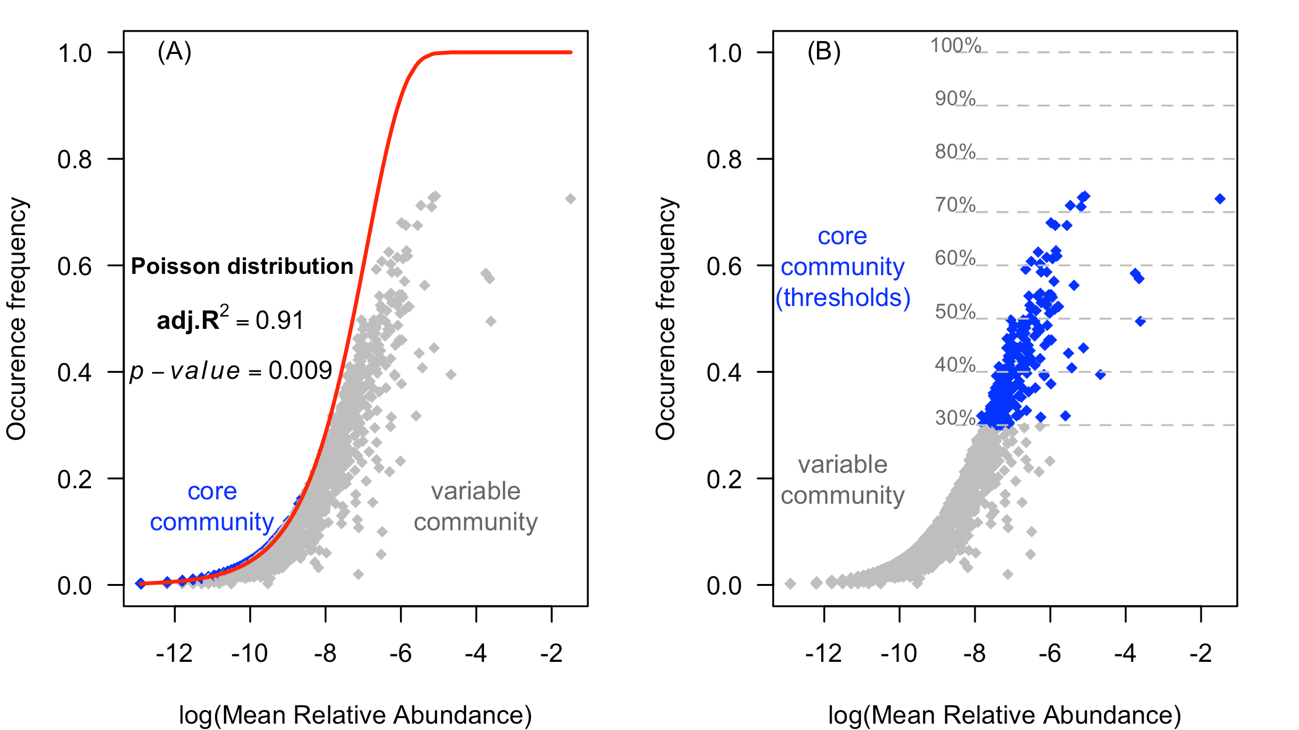

Figure S12 – The core and variable communities of the grape microbiome determined by (A) our proposed method based on the Poisson distribution and (B) an arbitrary, threshold-based method.

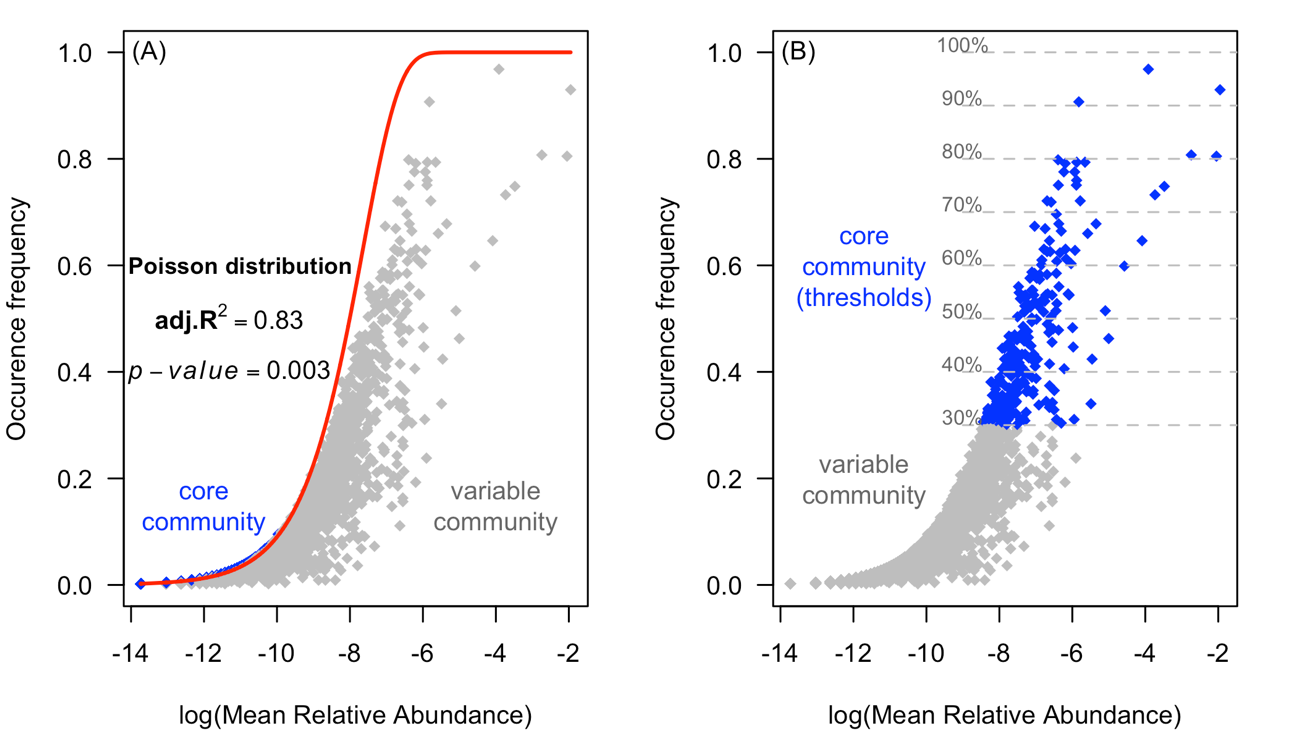

Figure S13 – The core and variable communities of the maize microbiome determined by (A) our proposed method based on the Poisson distribution and (B) an arbitrary, threshold-based method.

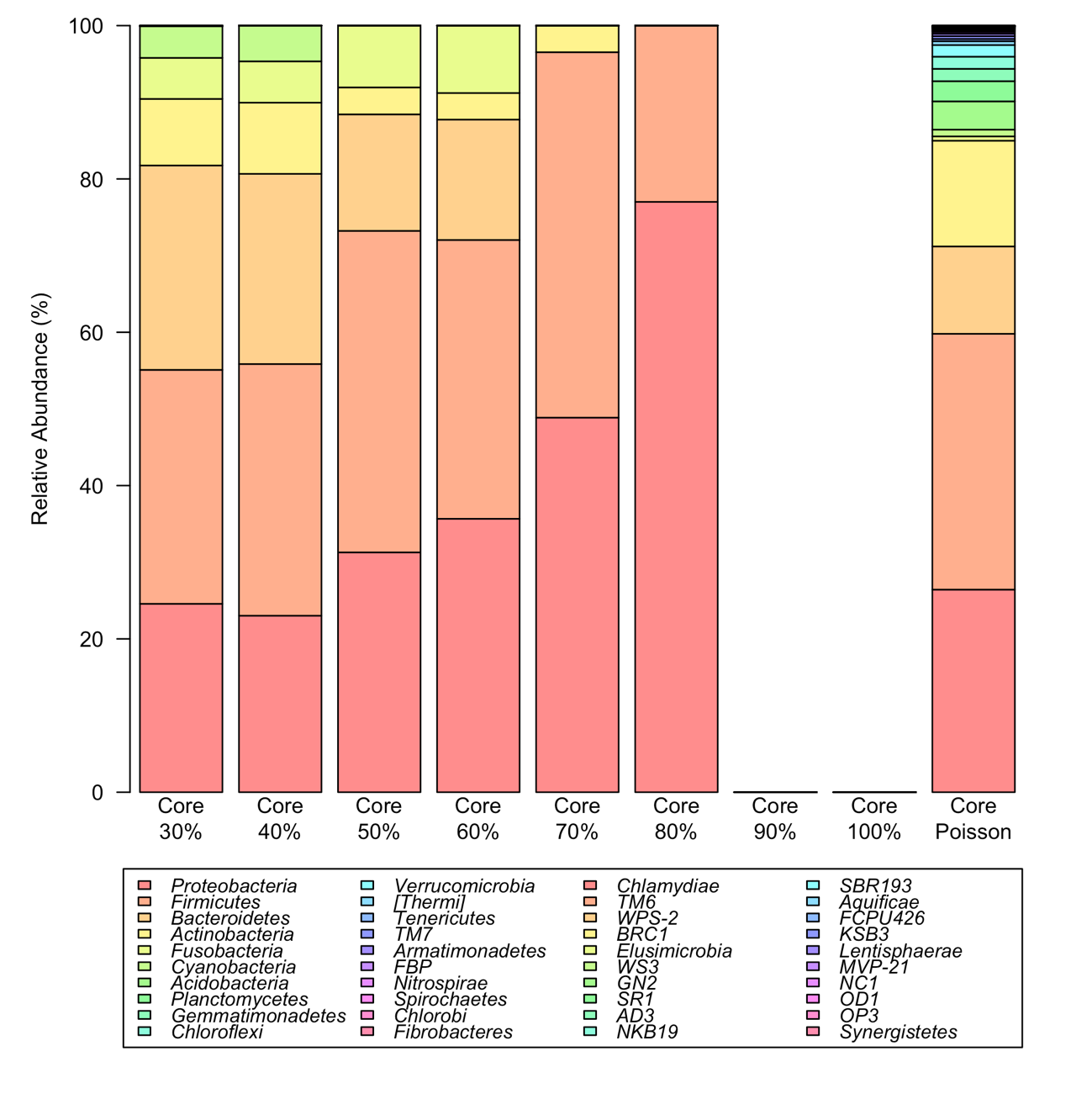

Figure S14 - Percentage of relative abundance of the core communities of human database determined by conventional methods (thresholds of 30,40,50,60,70,80,90 and 100%) and by Poisson distribution model (Core Poisson).

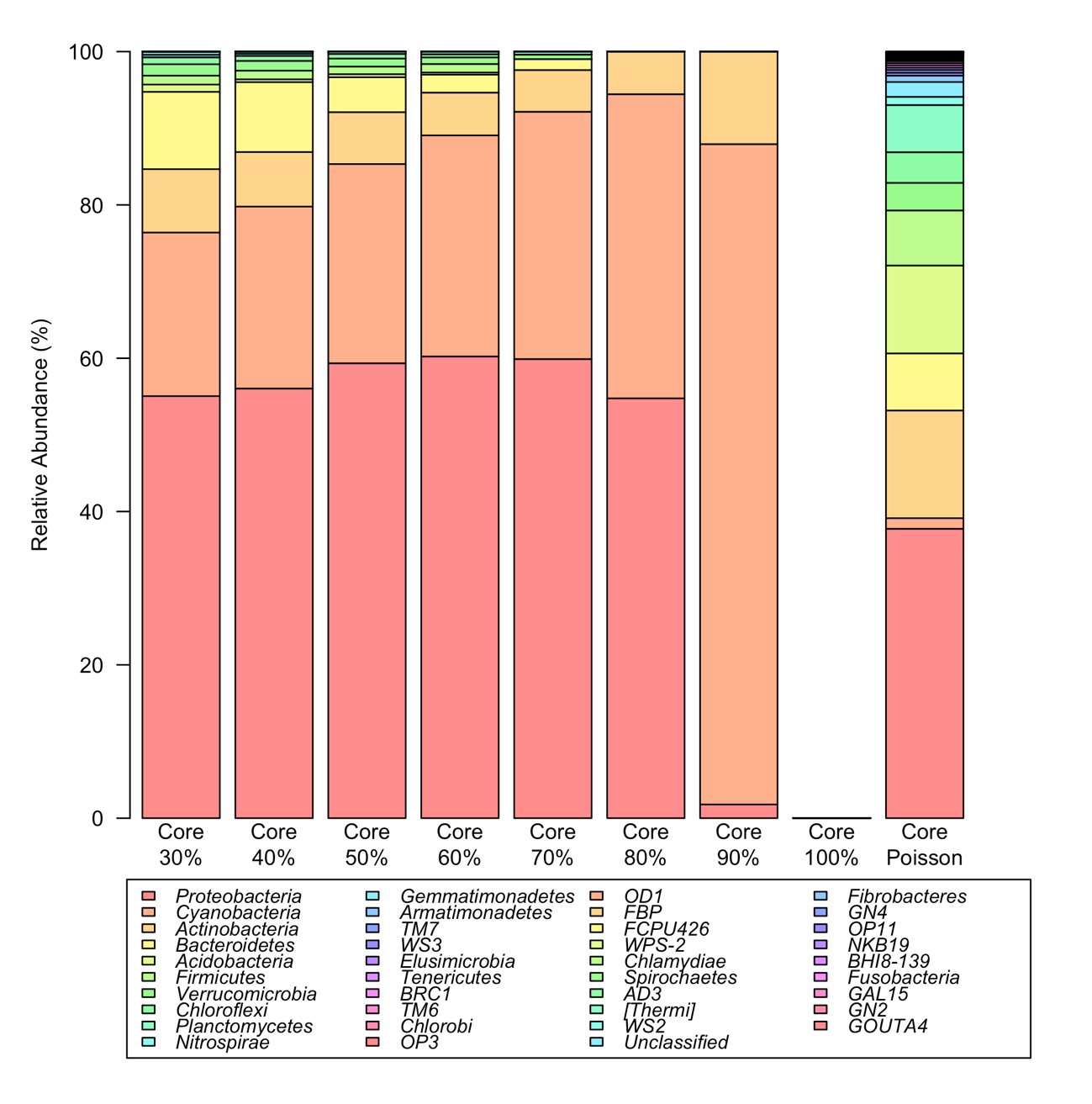

Figure S15 – Percentage of relative abundance of the core communities of maize database determined by conventional methods (thresholds of 30,40,50,60,70,80,90 and 100%) and by Poisson distribution model (Core Poisson).

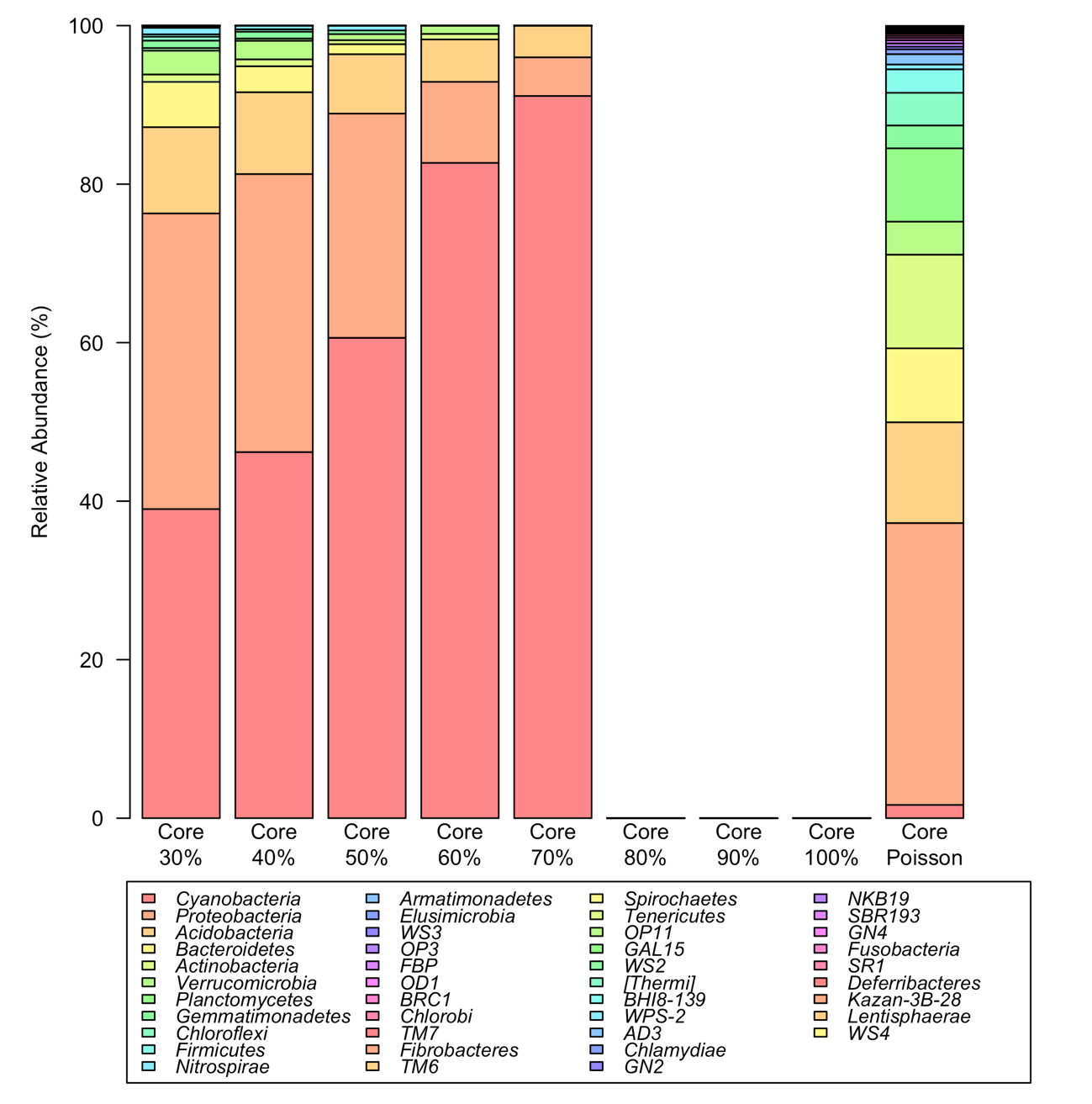

Figure S16 – Percentage of relative abundance of the core communities of grape database determined by conventional methods (thresholds of 30,40,50,60,70,80,90 and 100%) and by Poisson distribution model (Core Poisson).
